# *SPP1*^hi^ macrophages in fibrin niches promote hyperplastic tissue remodeling in rheumatoid arthritis synovium

**DOI:** 10.64898/2026.04.29.721703

**Authors:** Ian Mantel, Haoxuan Zhang, Juan Vargas, Gao Ce, Hope Townsend, Richard Bell, Amit Lakhanpal, Miriam R. Fein, Accelerating Medicines Partnership: RA/SLE Network, Thomas Norman, Dana Orange, Daniel Ramirez, Edward F. DiCarlo, Susan M. Goodman, Melanie H. Smith, Fan Zhang, Kevin Wei, Kushal K. Dey, Alexander Rudensky, Christina S. Leslie, Laura T. Donlin

**Author notes:** These authors contributed equally (Ian Mantel, Haoxuan Zhang).

## Abstract

In chronic inflammatory diseases, maladaptive tissue remodelling is driven by a complex interplay of resident cells, immune infiltrates and the extracellular matrix. In the autoimmune disorder rheumatoid arthritis (RA), synovial tissue undergoes assive expansion to form an invasive pannus that drives the erosion of cartilage and bone. The mechanisms mediating this aggressive growth are incompletely defined. Using spatial transcriptomics profiling of patient tissue, we detected an abundance of proliferating fibroblasts near the synovial tissue lining surface and adjacent to *SPP1*^hi^ macrophages. Notably, these synovial lining regions were also distinctly marked by deposits of the clot-forming protein fibrin. While the *SPP1*^hi^ acrophages phenotypically resemble pro-fibrotic macrophages that drive lung and liver fibrosis, these niches were devoid f the dense highly ordered collagen that marks fibrosis. Functionally, we found that *SPP1*^hi^ macrophages degrade and phagocytose fibrin matrices and promote fibroblast proliferation. As fibrin provides transient matrices for de novo tissue generation in the context of wound healing, these data support a model of hyperplastic tissue outgrowth involving *SPP1*^hi^ acrophages, fibroblasts and fibrin matrices adhered to the exterior synovial tissue surface. While current RA therapies primarily aim to dampen pro-inflammatory responses, our findings provide rationale for targeting pro-generative pathways nd *SPP1*^hi^ macrophages.

**Once Sentence Summary:** *SPP1*^hi^ macrophages in RA synovial fibrin deposits promote tissue hyperplasia.

## Introduction

Chronic inflammatory conditions commonly promote maladaptive tissue remodeling. Persistent inflammatory infiltrates deregulate and divert homeostatic and tissue repair processes, leading to a variety of pathological consequences. Macroscopically, tissues may expand, atrophy or become deformed, while the underlying interstitial matrix can stiffen or loosen. Concurrently, resident cells exhibit a range of responses, including cell death, excessive proliferation, transdifferentiation or senescence, while infiltrating immune cells stably engraft into the tissue architecture.

In rheumatoid arthritis (RA), the synovial tissue undergoes a profound transformation, expanding from a thin lining into a bulky, highly vascularized mass that can exceed 20 times its original volume (*1–3*). The synovium can also remodel into villous-like projections and pannus (*4*), an invasive tissue that erodes cartilage and bone. The mechanisms permitting this aggressive expansion remain poorly understood.

At the cellular level, RA synovium is marked by hyperplasia of resident fibroblasts and the infiltration of immune cells that form ectopic lymphoid structures. Recent studies have elucidated a complex cellular landscape, identifying more than 70 discrete cellular states (*5, 6*). However, it remains unclear which cells drive synovial tissue neogenesis and whether this occurs deep within the tissue or at the luminal surface.

Regenerative tissue responses, such as those involved in wound healing, require cooperation between stromal and blood-derived cells, including neutrophils, macrophages, fibroblasts, and endothelial cells, along with various extracellular matrix factors. Dysregulated regenerative networks result in divergent pathologic states, including fibrosis, where formation of stiff collagen deposits disrupts parenchymal tissue function. Notably, macrophages marked by high expression of *SPP1* are increasingly recognized as key contributors to fibrotic disorders across various tissues such as the lung and liver, where they associate with poor clinical outcomes (*7–16*). Although RA is widely described as an inflammatory disease driven by type 1 immunity and not a prototypical fibrotic disorder, RA synovial tissue frequently contains *SPP1*^hi^ macrophages, with roughly one third of patient tissue dominated by these cells (*5, 17–19*). It remains unclear if *SPP1*^hi^ macrophages in the RA synovium induce fibrosis or mediate some other form of tissue remodelling.

In this study, we investigated how RA synovial tissue undergoes massive expansion. Through spatial transcriptomic profiling of patient tissues and cell-based assays, we found proliferating and matrix remodeling fibroblasts near *SPP1*^hi^ macrophages, which were embedded in fibrin matrices that extended from the synovial lining surface. To functionally characterize this population, we established an in vitro system to recapitulate the *SPP1*^hi^ macrophage phenotype, which phagocytosed fibrin aggregates and supported fibroblast proliferation. Our data support a model of hyperplastic tissue remodeling at the lining surface in which fibrin deposits provide a transient matrix for de novo synovial tissue generation mediated by *SPP1*^hi^ macrophages and fibroblasts.

### *SPP1*^hi^ macrophages occupy an RA synovial lining niche with fibroblasts primed for proliferation and matrix deposition

To gain insights into the cellular and molecular features underlying RA-associated synovial tissue outgrowth, we profiled 10 RA patient synovial tissue samples using the Xenium spatial transcriptomics platform. The tissues were labeled with a 480-gene custom panel designed to distinguish broad cell lineages, RA synovial cell states (*5*), and pathways involved in tissue remodeling (**Supplementary Table 1A**). Tissues were selected from patients who met established diagnostic criteria and were positive for disease-associated anti-cyclic citrullinated peptide (CCP) and/or rheumatoid factor (RF) autoantibodies, and had a wide range of scores for histological metrics, including lymphocyte infiltration and lining hyperplasia (**Supplementary Table 1B**).

We used the Baysor cell segmentation algorithm (*20*) to identify 4,274,575 cells and assign them into one of six broad cell lineages (B cell, T cell, NK cell, myeloid, fibroblast, or endothelial) using SingleR (*21*) and an RA synovial cell reference dataset (*5*) (**Figure 1A, Supplementary Figure 1A**). The cells in each lineage were then further subclustered into a total of 33 granular cell states (**Supplementary Figures 1B-I, Supplementary Table 1C-J**). We next performed niche clustering based on the relative cell type composition of local spatial neighborhoods (**Methods**) and identified five niches shared across the 10 tissues (**Figure 1B**). Niches 1 and 5 contained the highest proportion of lymphocytes and were typically spatially adjacent to each other. Plasma cell aggregates were most prominent in Niche 1, while Niche 5 contained CD4^+^ T cells (T0) and B cells (B1) that constituted ectopic lymphoid structures (**Figure 1C-D; Supplementary Table 1K; Supplementary Figure 1J-K**). Niche 2 occupied a perivascular location and largely consisted of endothelial (E1-4) and mural (F3) cells. Niche 3 visually marked the synovial lining and contained the highest proportion of *HBEGF*^+^ lining fibroblasts (F2) and *SPP1*^hi^ macrophages (M2). Niche 4 encompassed interstitial regions, with sublining fibroblasts marked by *FNDC1* (F0) and *SFRP1* (F1) as the key cellular constituents.

**Figure 1.**
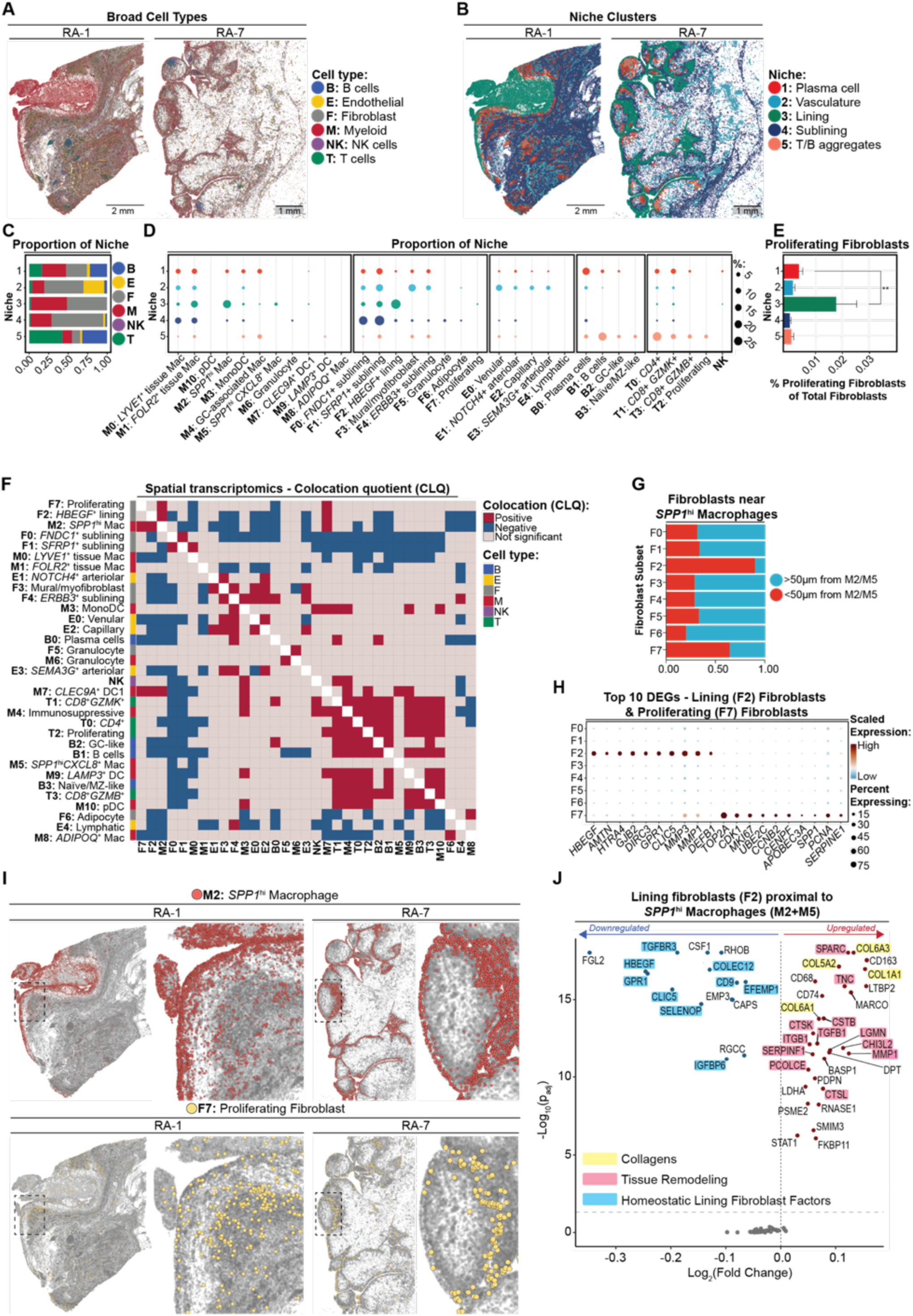
*SPP1*^hi^ macrophages occupy an RA synovial lining niche with fibroblasts primed for proliferation and matrix deposition. **A.** Spatial transcriptomics images of RA synovial tissue with the cells segmented and colored based on their broad cell lineage assignments. n=10 patient samples profiled, two representatives shown. F (grey)=fibroblasts; M (red)=myeloid cells; T (green)=T cells; B (blue)=B cells; E (yellow)=endothelial cells; NK (purple)=natural killer cells. **B.** Spatial transcriptomics images with cells colored based on Niche assignment. Niche 1 (red)=Plasma cell; Niche 2 (blue)=Vasculature; Niche 3 (green)=Lining; Niche 4 (navy)=Sublining; Niche 5 (salmon)=T/B aggregates. **C.** The proportion of the broad cell types in each Niche across the n = 10 tissues **D**. The distribution of each fine cell cluster across the 5 Niches across the n = 10 tissues. Dot size corresponds to the proportion in each Niche. **E**. The proportion of proliferating fibroblasts (F7) of total fibroblasts in each Niche across the n = 10 tissues. **Wilcoxon matched-pairs signed rank test, *p* = 0.0039. **F.** Colocation quotient (CLQ) matrix for each cell subset, with the query cell represented on the y-axis; significance cutoff <0.05 (BH corrected across cell type pairs and samples). **G.** The proportion of each fibroblast subtype that is <50µm (red) or >50µm (blue) from an *SPP1*^hi^ macrophage (either an M2 or M5 cell) across the n = 10 tissues. **H.** Dot plot heatmap displaying how the fine clusters of fibroblasts express the top 10 differentially expressed genes (DEGs) between lining fibroblasts (F2) and proliferating fibroblasts (F7) compared to all cells. **I.** Spatial transcriptomic images with cells represented by colored dots for either *SPP1*^hi^ macrophages (M2) (red) or proliferating fibroblasts (F7) (yellow). Region of interest marked by dotted line is magnified on the righthand image. **J.** Volcano plot displaying Niche-DE analysis of lining fibroblasts (F2) gene expression changes based on proximity to *SPP1*^hi^ macrophages (M2+M5).

To identify areas of putative active tissue growth, we first quantified the proportion of proliferating fibroblasts (F7) within each niche. These proliferating fibroblasts distinctly expressed cell division related genes such as *CCNB2*, *CENPF*, *MKI67*, and *CDK1*; canonical lining fibroblast markers such as *MMP1*, *MMP3*, *PDPN*, and *FN1*; and activated lining fibroblast genes such as *COL5A2*, *FAP*, *TNC*, and *MT1G* (**Supplementary Table 1C-D,1H**) (*5, 22*). While Niches 2, 3, and 4 contained the highest overall proportions of fibroblasts, the synovial lining Niche 3 exhibited the greatest proportion of proliferating fibroblasts across the 10 RA tissues (**Figure 1C-E, Supplementary Table 1K;Supplementary Figure 1J-K**).

Next, independent of niche designation, we quantified the spatial enrichment of each cell state within 50µm of a target cell state using the colocation quotient (CLQ), which captures asymmetric spatial enrichment rather than mutual co-occurrence. We found that proliferating fibroblasts (F7) predominantly localized near two cell clusters: the *SPP1*^hi^ macrophages (M2) and the *CLEC9A*^+^ conventional type 1 dendritic cells (DC1) (M7) (**Figure 1F**). Given that the *SPP1*^hi^ macrophage (M2) cluster represented a more abundant cell type and the majority of these cells were localized in Niche 3 with high concentrations of proliferating fibroblasts, we focused subsequent analyses on this subset. As suggested by the CLQ analysis, a direct assessment across all fibroblast subsets demonstrated lining fibroblasts (F2) and proliferating fibroblasts (F7) had the highest proportions within 50µm of SPP1^hi^ macrophages (M2 + M5 clusters) (**Figure 1G, Supplementary Figure 1K**). Furthermore, we observed dense patches of proliferating fibroblasts (F7) spatially overlapping or directly adjacent to aggregates of *SPP1*^hi^ macrophages (M2) (**Figure 1I, Supplementary Table 1L**).

Using weighted gene expression analysis, we found the previously defined RA synovial *SPP1*^hi^ macrophage state identified by scRNA-seq on 79 patient samples (*5*) overlapped with our spatial transcriptomics-defined *SPP1*^hi^ macrophage (M2) cluster (**Supplementary Table 1M**, **Supplementary Figure 1L**). These subsets shared distinctively high levels of *SPP1*, *CLEC5A*, *AQP9*, and *FAPB5* (**Supplementary Tables 1C-D, 1J**), co-expression of markers like *CCR2* and *MARCO* that typically segregate monocytes and macrophages, respectively, and anti-inflammatory markers such as *TREM2* and *RNASE1.* In a minority of tissues profiled by spatial transcriptomics, we found a second less abundant *SPP1*^hi^ macrophage population (*SPP1*^hi^*CXCL8*^+^ (M5) cluster), which relative to the M2 *SPP1*^hi^ cluster, expressed select inflammatory markers (**Supplementary Table 1K**).

In a niche differential expression (Niche-DE) (*23*) analysis, we notably observed as the density of neighboring *SPP1*^hi^ macrophages increased, lining fibroblasts had higher expression of hallmark tissue generative factors including *COL1A1*, *COL5A2*, *COL6A1*, *COL6A3, TGFB1*, *TNC* (*24, 25*), which are otherwise predominantly expressed by sublining fibroblasts (**Figure 1J, Supplementary Table 1N**) (*5, 22, 26, 27*). These lining fibroblasts also increased canonical fibroblast extracellular matrix or tissue remodeling factors such as the integrin *ITGB1* that binds fibronectin; the proteases *CTSB*, *CTSK*, *CTSL, LGMN* (*28–31*); and the pro-collagen endopeptidase *PCOLCE* (*32*), as well as the lining fibroblast-defined factors *MMP1* and *MMP3*. Conversely, the more *SPP1*^hi^ macrophages that surrounded lining fibroblasts, the lower the expression of lining homeostatic markers, such as for *CLIC5, HBEGF, GPR1*, and *CD9* (*22*). This matrix remodeling signature also marked the proliferating fibroblast population (F7) (**Figure 1I-J, Supplementary Table 1C-D, 1H**). We next performed protein-protein interaction analysis on the 12 genes that were consistently upregulated in lining fibroblasts (F2) within 50µm of an SPP1^hi^ macrophage (in at least 8 of 10 tissues). This analysis showed that the genes were significantly interconnected (*p*=1.16 x 10^-4^), indicating these proteins form a functional module with key nodes in genes associated with extracellular matrix remodeling (*COL1A1*, *SPARC*) (**Supplementary Figure 1M**).Together, these data suggest that in synovial lining affected by RA, *SPP1*^hi^ macrophages coincide with activated lining fibroblasts with tissue generative programs poised for proliferation, collagen production and other key drivers of tissue growth.

### Synovial *SPP1*^hi^ macrophages colocalize with fibrin in a niche lacking dense collagen

Across the 10 RA samples, *SPP1*^hi^ macrophages (M2 and M5 clusters) largely formed dense aggregates, either in a linear array at the synovial lining or as more substantial multi-layered clusters below the lining **(Figure 2A, Supplementary Figure 1K).** By contrast, other myeloid cells such as those in the Monocyte-Dendritic Cell (M3: MonoDC) cluster were relatively evenly distributed throughout tissue compartments and lacked evidence of spatial aggregation (**Supplementary Figure 1E**). To gain insights into the extracellular matrix composition of regions occupied by *SPP1*^hi^ macrophages, we first stained tissue sections with hematoxylin and eosin (H&E) and imaged by light microscopy (**Supplementary Figure 2A**). We then binarized each tissue into regions defined by the presence (≥1 cell) or absence of *SPP1*^hi^ macrophages and overlaid outlines of these regions atop the corresponding H&E images (**Supplementary Figure 2B-C**). Compared to the long, parallel, undulating collagen fibers of canonical synovial interstitial tissue, the extracellular matrix in *SPP1*^hi^ macrophage-containing tissue regions appeared amorphous, with bright eosin staining. These histologic features of *SPP1*^hi^ macrophage-containing regions were consistent with fibrin, a transient sealant and scaffolding matrix important in wound repair and blood clots (**Figure 2B**, **Supplementary Figure 2B**) (*33*).

**Figure 2.**
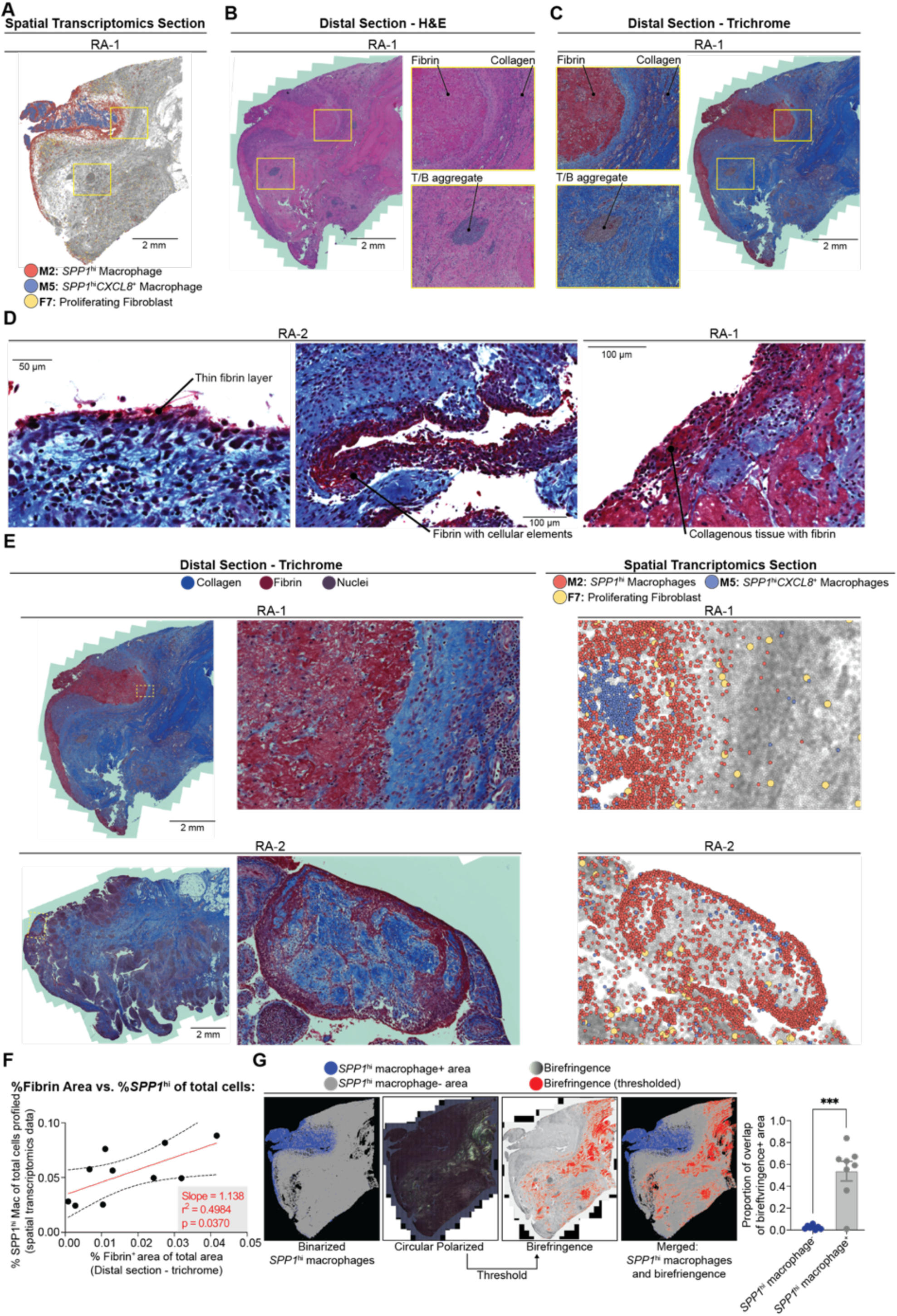
Synovial *SPP1*^hi^ macrophages colocalize with fibrin in a niche lacking dense collagen. **A.** Representative RA patient synovial tissue (n=10) spatial transcriptomics image depicting *SPP1*^hi^ macrophages (M2, red), *SPP1*^hi^*CXCL8*^+^ macrophages (M5, blue) and proliferating fibroblasts (F7, yellow). n=10. **B-C.** Images of distal sections for the tissue in panel A stained with H&E (**B**) and Masson’s Trichrome (**C**); regions of interest (dashed yellow boxes) shown in higher magnification off to the right (B) or left (C). **D.** Selected images of Trichrome stained regions depicting different patterns of fibrin staining. **E.** Representative images of Trichrome stained tissue from a distal section (left, center), and spatial transcriptomics-labeled section (right). RA-2 region of interest at higher magnification has been rotated clockwise ∼90 degrees. **F.** Correlation plot of the percentage of tissue stained by Trichrome and identified as a fibrin deposit (x-axis) and the proportion of *SPP1*^hi^ macrophages of total cells as determined by spatial transcriptomics analysis (y-axis) across the n = 10 tissues; red line represents simple linear regression fit, with *F*(1,8) = 6.25, *p* = 0.04; dashed lines represent 95% CI. **G.** Representative images (from left): tissue binarized into two regions depending on the presence or absence of *SPP1*^hi^ macrophages; birefringence signal from circularly polarized light microscopy; thresholded birefringence signal; merging of birefringence and binarized images; summary of calculation of the extent of overlap between *SPP1*^hi^ containing regions and birefringence regions. Across n = 8 tissues with intact Xenium slides after analysis and transport; ***Paired t-test, *p* = 0.0006.

To validate these findings, we stained three additional sections from each tissue with a panel of histologic dyes: Masson’s Trichrome, which stains fibrin in red and collagen in blue; phosphotungstic acid-hematoxylin (PTAH), which visualizes fibrin in blue and collagen and nuclei in purple; and H&E, which stains fibrin bright pink and collagen pale pink (**Figure 2B-C, Supplemental Figure 2D-F**). Collectively, these stains confirmed that fibrin accumulated in thin layers at specific sites in the synovial lining, but also formed massive embedded aggregates as large as three millimeters wide in several tissues **(Figure 2C)**. High magnifications revealed fibrin organized into a glassy reticular network, permeating the interstitium around cells (**Figure 2D, Supplementary Figure 2A, D-F**). Notably, in regions with preserved architecture across the sections, we observed a consistent colocalization of fibrin-rich regions and *SPP1*^hi^ macrophages (**Figure 2E**). Accordingly, using an object classification algorithm, we found a positive correlation between the percentage of area containing fibrin in each RA sample and both the proportion and the total area percentage of *SPP1*^hi^ macrophages (M2+M5) (**Figure 2F, Supplementary Figure 2G, Supplemental Table 2A, Methods**).

As fibrin provides a provisional matrix in tissue repair, the association of *SPP1*^hi^ macrophages within fibrin deposits suggested these cells accumulated in regions undergoing tissue remodeling. To gain further insights into the surrounding tissue properties, we reimaged with circularly polarized light microscopy and measured birefringence. In fibrotic tissue and scars, dense, highly ordered collagen fibers generate high birefringence signals. Upon overlaying these signals onto the spatial transcriptomic images, we found the *SPP1*^hi^ macrophage-containing and surrounding regions were devoid of high birefringence (**Figure 2G, Supplementary Figure 2H, Supplementary Table 2B**). Thus, the RA synovial lining regions containing and immediately surrounding fibrin and *SPP1*^hi^ macrophages were notably absent of dense collagen fibers that mark fibrosis.

### RA synovial *SPP1*^hi^ macrophages exhibit pro-fibrotic macrophage features

SPP1-expressing (*SPP1*^hi^) macrophages drive tissue remodeling in fibrosis. To understand their function in the rheumatoid arthritis (RA) synovium, we compared them with their counterparts from a fibrotic lung and liver scRNA-seq atlas. Reference mapping showed that 67% of pro-fibrotic *SPP1*^hi^ macrophages from lung and liver aligned with RA synovial *SPP1*^hi^ macrophages (**Figure 3A, Supplementary Table 3A, Supplementary Figure 3A**). Furthermore, top marker genes for fibrotic tissue *SPP1*^hi^ macrophages, such as *SPP1*, *CD9*, *LGALS3*, and *FABP5*, were also highly expressed in the RA synovial *SPP1*^hi^ subset (**Figure 3B**). These findings highlight a conserved transcriptional signature for *SPP1*^hi^ macrophages across diverse diseases and tissues.

**Figure 3:**
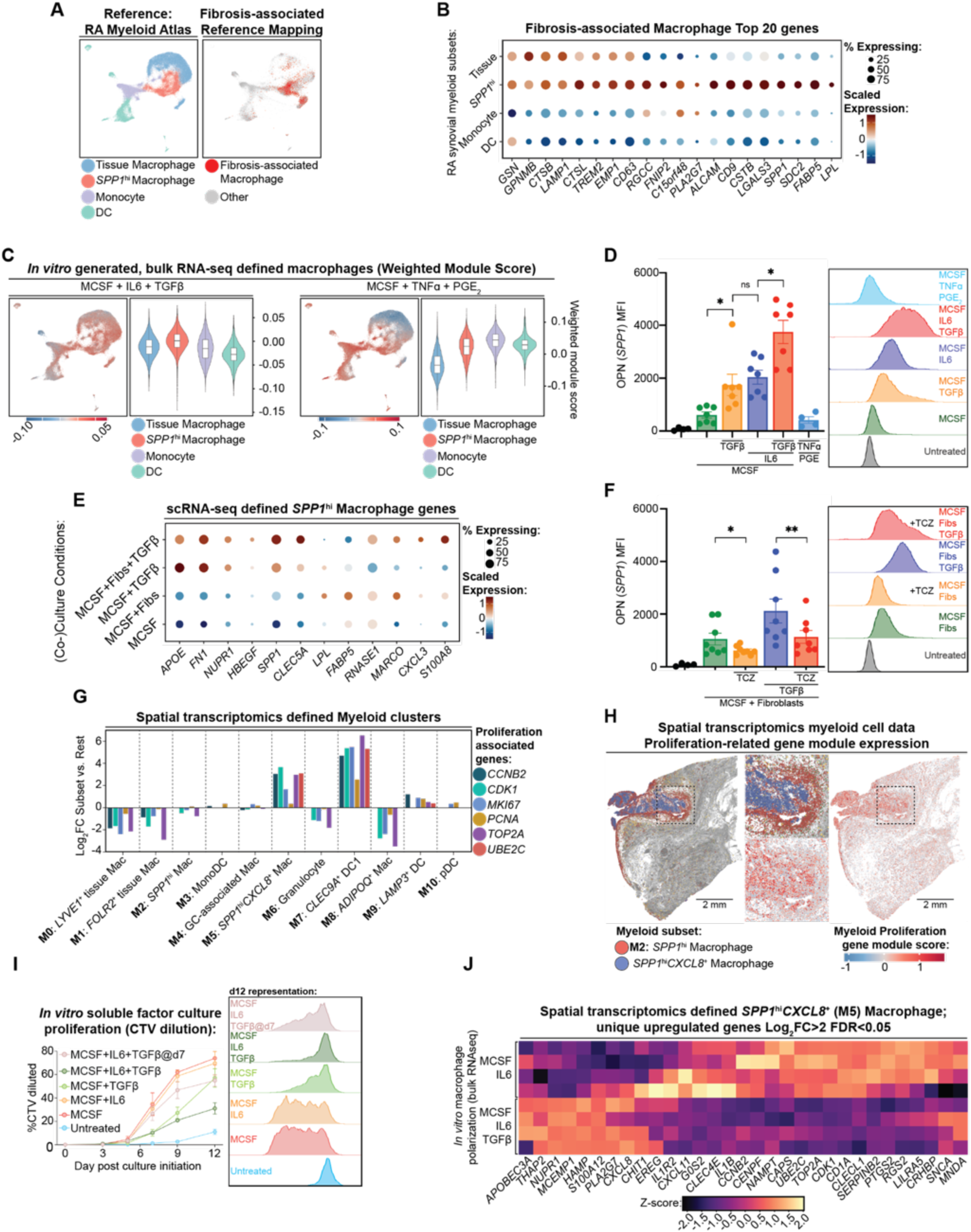
R**A synovial macrophages exhibit pro-fibrotic macrophage features. A.** Reference mapping of fibrosis-associated macrophages from lung and liver(*8*) (red) onto RA synovial myeloid scRNAseq reference data and UMAP(*5*). **B.** Dotplot expression levels for the top 20 fibrosis-associated macrophage markers expressed by RA synovial myeloid subsets. **C.** Weighted module score for differentially expressed genes from bulk RNAseq of human CD14^+^ monocytes treated for 72h with M-CSF, TNF-ɑ, and PGE2 (left side) or M-CSF, IL-6, and TGF-β (right side) compared to M-CSF alone. The score of each cell in UMAP corresponds to legend under; violin plot shows weighted module score for each cell within indicated subset. For violin plots, One-sided Wilcoxon Rank Sum Test of all myeloid subsets vs. *SPP1*^hi^ macrophage p<0.0001 **D.** Intracellular flow cytometry detection of *SPP1*-encoded osteopontin (OPN) in CD14^+^ monocytes treated for 72h. n = 8. Wilcoxon signed rank test; *M-CSF vs. M-CSF and TGF-β, *p* = 0.016; *M-CSF and IL-6 vs. M-CSF, IL-6 and TGF-β, *p* = 0.016. **E.** Dotplot expression levels of scRNAseq-defined *SPP1*^hi^ macrophage cluster marker genes in macrophages CD14^+^ cells treated with M-CSF for 24h) (co-)cultured as indicated for 72h. **F.** Intracellular flow cytometry analysis for OPN levels in monocytes treated and exposed to synovial fibroblasts as indicated for 72h with or without tocilizumab (TCZ). n = 8. Wilcoxon signed rank test; *M-CSF+Fibs vs. M-CSF+Fibs+TCZ, *p* = 0.016; **M-CSF+Fibs+TGF-β vs. M-CSF+Fibs+TGF-β+TCZ, *p* = 0.008. **G.** Spatial transcriptomics expression log2 fold-change (FC) gene expression changes for each Myeloid subset compared to all other cells. **H.** Representative spatial transcriptomic images with location of *SPP1*^hi^ macrophages (M2, red) and *SPP1*^hi^*CXCL8*^+^ macrophages (M5, blue) (left) and each myeloid cell scored for the level of a proliferation gene expression module (comprised of genes shown in (**I**)) (right). Module scores were capped at the 99.5th percentile to improve contrast on scale. **I.** Proliferation assay time course summary (left) and day 12 flow cytometry histogram (right) for Cell Trace Violet (CTV) dilution in monocytes treated with indicated factors for 12 days; (MCSF+IL6+TGFβ@d7 represents M-CSF and IL-6 for 7 days followed by TGF-β added on day 7). **J.** Heatmap of expression of genes uniquely expressed by *SPP1*^hi^*CXCL8*^+^ macrophages (M5) (defined as log2FC>2, FDR<0.05 in M5, not in M2).

To identify signaling pathways driving the *SPP1*^hi^ phenotype in RA, we turned to factors known to generate these macrophages in fibrosis. Previous studies established that a combination of three mediator categories is required: 1) a macrophage growth factor (e.g., M-CSF, GM-CSF), 2) a STAT3-activating cytokine (e.g., IL-6, IL-17), and 3) a third factor, such as TGF-β (*8, 13, 34*). Guided by this framework, we selected a pro-fibrotic cocktail using M-CSF, due to its high expression by synovial fibroblasts compared to GM-CSF (**Supplementary Figure 5B**)(*5*); IL-6, given the clinical efficacy of IL-6R inhibitors in RA(*35*) and its high expression by synovial fibroblasts (*36, 37*); and TGF-β, for its central role in tissue remodeling (*5, 8, 38, 39*). We contrasted this with a pro-inflammatory condition (M-CSF, TNF-ɑ, PGE2) that we have shown induces an inflammatory macrophage phenotype (*IL1B*^+^*HBEGF*^+^) (*40*). Bulk RNA sequencing revealed that the gene module induced by the pro-fibrotic cocktail was highly expressed in the synovial *SPP1*^hi^ macrophage subset, while the pro-inflammatory cocktail aligned with inflammatory monocytes (**Figure 3C, Supplementary Figure 3C-D, Supplementary Table 3B**). At the protein level, the pro-fibrotic combination synergistically increased osteopontin (encoded by *SPP1*), whereas the pro-inflammatory condition did not (**Figure 3D**). These results demonstrate that M-CSF, IL-6, and TGF-β induce a macrophage state that closely resembles RA synovial *SPP1*^hi^ macrophages.

Given the abundance of fibroblasts in the synovial lining (Niche 3), we tested their influence on the *SPP1*^hi^ phenotype. We co-cultured M-CSF-pretreated monocytes with synovial fibroblasts and TGF-β. This combination induced the highest proportion of cells matching the RA synovial *SPP1*^hi^ macrophage phenotype and expressing key marker genes (**Figure 3E, Supplementary Figure 3E, Supplementary Table 3C**). Furthermore, fibroblasts plus TGF-β induced osteopontin production, an effect blocked by the IL-6R inhibitor tocilizumab (**Figure 3F**). This suggests synovial fibroblasts promote the *SPP1*^hi^ macrophage state via IL-6 signaling.

Lastly, we investigated the proliferative potential of *SPP1*^hi^ macrophages. Spatial transcriptomics revealed a proliferative signature (e.g., *MKI67*, *TOP2A*) in a specific *SPP1*^hi^*CXCL8*^hi^ macrophage subset located within fibrin deposits (**Figure 3G, 3H**). To explore this in vitro, we tested if the pro-fibrotic factors induced cell division. We found that M-CSF drove proliferation regardless of the presence of IL-6, while TGF-β delayed its onset (**Figure 3I**). The discrepancy between the in vivo signature and in vitro results suggests that the unique microenvironment of fibrin deposits, where the proliferative *SPP1*^hi^*CXCL8*^+^ subset resides, may provide specific temporal or combinatorial signals that promote cell division (**Figure 3J, Supplementary Figure 3F**).

Collectively, these observations indicate a high degree of conservation between the *SPP1*^hi^ macrophage phenotype found in fibrosis and the RA synovium and further suggest overlapping signaling pathways that differentiate these macrophage states across tissues and diseases.

### *SPP1*^hi^ macrophages digest fibrin and support fibroblast proliferation

Given their striking overlap with fibrin deposits and a gene signature for clot formation and coagulation, including high levels of *SERPINE1* and *F13A1,* we initially hypothesized that RA synovial *SPP1*^hi^ macrophages would induce fibrin polymerization to form a clot. To assess this potential, we incubated activated macrophage subsets with human plasma and measured how turbidity changed over time. Inflammatory macrophages, differentiated by M-CSF and TNF-ɑ with or without PGE2, rapidly promoted clotting, while cells polarized by M-CSF with or without IL-6 induced clotting at a slower rate. Conversely, the three-factor in vitro-derived *SPP1*^hi^ macrophages did not promote clotting and prevented the low level of spontaneous clotting observed in cell free plasma (**Figure 4A**). This suggested that while *SPP1*^hi^ macrophages colocalize with fibrin deposits, they likely do not contribute to clot formation.

**Figure 4.**
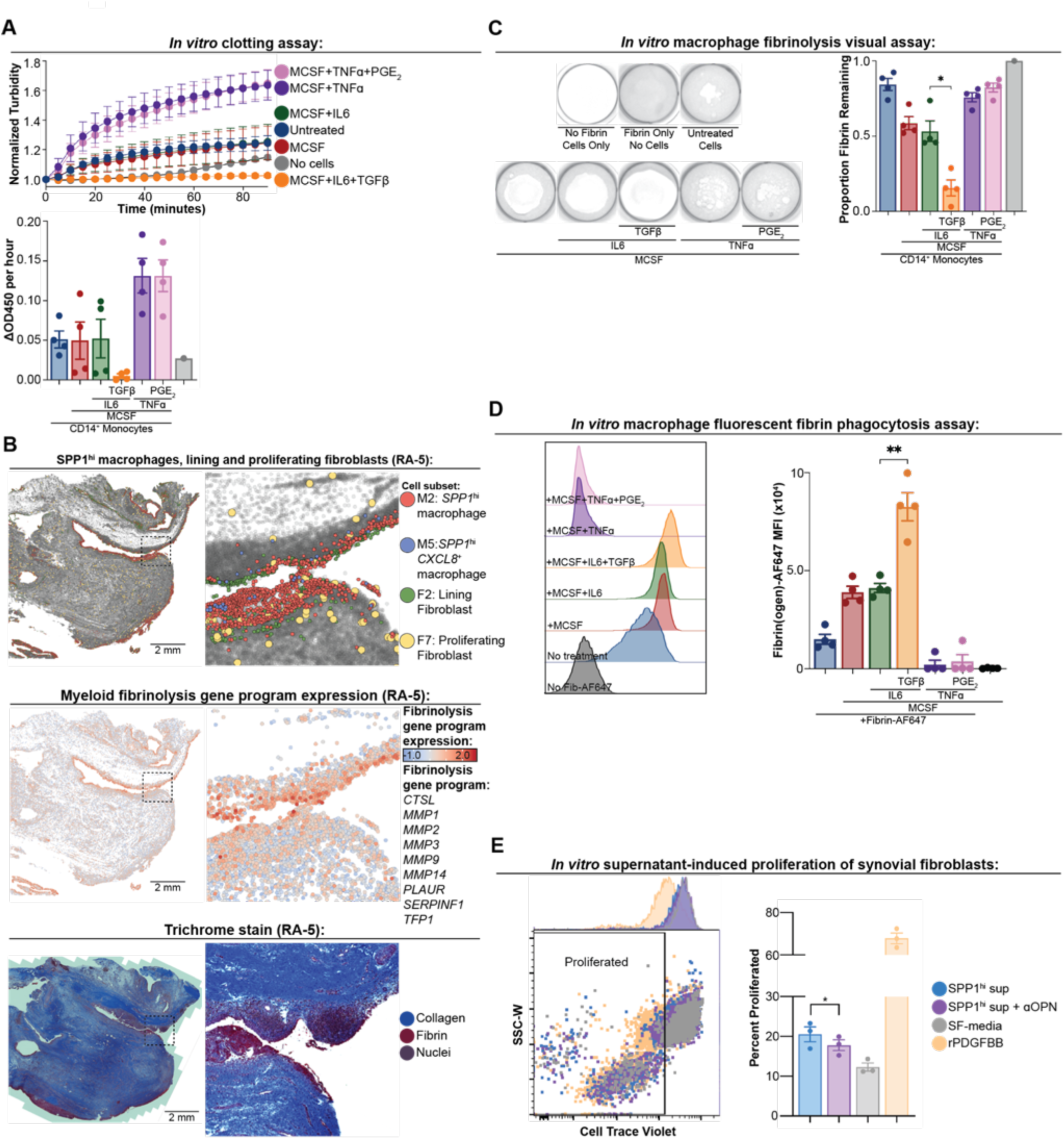
*SPP1*^hi^ macrophages digest fibrin deposits and support fibroblast proliferation. **A.** Plasma clotting assay measuring changes in turbidity of human plasma clotting assay after exposure to CD14^+^ monocytes pre-polarized for 72h with the indicated soluble factors, with the time course (top) and fold-change over 72h (bottom). n=4 biologic replicates. **B.** Representative spatial transcriptomics/histological images of: (top panel) *SPP1*^hi^ macrophages (M2, red), *SPP1*^hi^CXCL8^+^ macrophages (M5, blue), lining fibroblasts (F2, green), and proliferating fibroblasts (F7, yellow); (middle panel) score for the fibrinolysis-associated gene program in myeloid cells; (bottom panel) Masson’s trichrome stain for a distal section. **C.** Representative images of fibrinolysis assay with fibrin gels and CD14^+^ monocytes exposed to indicated soluble factors for 6d (left) and with the Coomassie-stained fibrin levels remaining at day 6 plotted (right). n = 4 biological replicates. Paired t-test; M-CSF+IL-6 vs. M-CSF+IL-6+TGF-β. *Paired t-test, *p* = 0.022. **D.** Representative result of in vitro fluorescent fibrin phagocytosis flow cytometric assay from monocytes pre-polarized with indicated factors and incubated atop fibrin gels containing AF647-labelled fibrinogen for 24h (left) and mean fluorescent intensity (MFI) plotted (right); n = 4 biological replicates; **Paired t-test; M-CSF+IL-6 vs. M-CSF+IL-6+TGF-β *p =* 0.004. **E.** Proliferation assay with Cell Trace Violet (CTV) dilution for synovial fibroblasts treated with indicated in vitro generated monocyte-derived supernatant/soluble factors. *Paired t-test; *SPP1*^hi^ sup vs. *SPP1*^hi^ sup + ɑOPN *p* = 0.035.

Many gene products associated with coagulation and clot formation pathways are involved in the breakdown of fibrin matrices referred to as fibrinolysis. These genes, which include *MMP1*, *MMP3*, *CTSL*, and *PLAUR*, are highly expressed by RA synovial *SPP1*^hi^ macrophages along with synovial fibroblasts. Notably, we observed a colocalization of myeloid cells with high expression of a fibrinolysis gene program and *SPP1*^hi^ macrophages, particularly at the very edge of the tissue where fibrin deposits are observed (**Figure 4B**). To explore macrophage fibrinolysis in vitro, CD14^+^ cells were differentiated on fibrin gels for 5 days. We observed that in wells containing cells differentiated with M-CSF, IL-6 and TGF-β, which induces the *SPP1*^hi^ macrophage phenotype (**Figure 3D-F**), the fibrin gel had completely dissolved by day 5. By contrast, CD14^+^ cells cultured with pro-inflammatory factors had a nearly intact fibrin gel. Following up on this observation, we next developed an in vitro fibrin digestion visual assay, which confirmed that *SPP1*^hi^ macrophages possess a unique ability to dissolve fibrin upon which they are cultured (**Figure 4E**). While these cells actively degraded fibrin matrices, we found no change in the core *SPP1*^hi^ macrophage phenotype after fibrin exposure by RT-qPCR or flow cytometry **(Supplementary Figure 4A-B**).

Consistent with their superior degradative capacity, *SPP1*^hi^ macrophages also exhibit upregulated phagocytosis associated genes like *MARCO* and *TREM2*, both of which are involved in the clearance of cellular debris (**Supplementary Table 1D, 1J**). Given their superior ability to degrade fibrin over the course of culture, we next assessed the ability of different macrophages to engulf fibrin. We cultured pre-polarized macrophages atop a fibrin gel containing fluorescently-labelled fibrinogen for 24h, then analyzed the cells by flow cytometry. This assay revealed that *SPP1*^hi^ macrophages engulfed the highest amount of fibrin, while the pro-inflammatory macrophage subset engulfed no detectable levels (**Figure 4D**).

In addition to localization to fibrin deposits, our spatial transcriptomics also revealed a highly specific colocalization between *SPP1*^hi^ macrophages and proliferating fibroblasts (F7). In an in vitro co-culture assay with a blocking antibody to the *SPP1*-encoded protein osteopontin, we found *SPP1*^hi^ macrophages supported fibroblast proliferation via secreted osteopontin (**Figure 4E**).

Taken together, these results demonstrate that *SPP1*^hi^ macrophages perform two key functions in the RA synovial niche: they support fibroblast proliferation in an osteopontin-dependent manner and they actively contribute to fibrin polymer degradation. This latter process is perhaps mediated by MMP and cathepsin-dependent enzymatic fibrinolysis and, ultimately, internalization of fibrin breakdown products.

### *SPP1*^hi^ macrophage targeting by IL-6 receptor inhibition in RA synovium

Based on the reported efficacy of the IL-6 receptor inhibitor in patients with Myeloid-rich synovium (*35*), we hypothesized this clinical response was mediated in part by targeting of *SPP1*^hi^ macrophages given that our prior collaborative study found *SPP1*^hi^ macrophages dominate Myeloid-rich tissues (*5*). To test this, we first performed a Weighted Gene Co-Expression Network Analysis (WGCNA) on the synovial biopsy bulk RNA-seq data from the tocilizumab and rituximab (a B cell targeting therapy) treatment arms of the aforementioned clinical study (*35*). Paired differential analysis identified two dominant modules repressed by the medications (Module 2 and 4). Both medications repressed Module 4, which characterized as a broad inflammatory and immune signaling pathway in an Over-Representation Analysis. In contrast, tocilizumab uniquely repressed Module 2, which was enriched for fundamental biosynthetic and metabolic processes (**Figure 5A, Supplementary Table 5A**). Spatial mapping of the Module 2 genes showed enrichment in the Synovial Lining Niche (Niche 3), potentially linking this tocilizumab-specific effect to the spatial location of *SPP1*^hi^ macrophages, fibrin and proliferating and matrix remodeling fibroblasts (**Figure 5B-C**).

**Figure 5.**
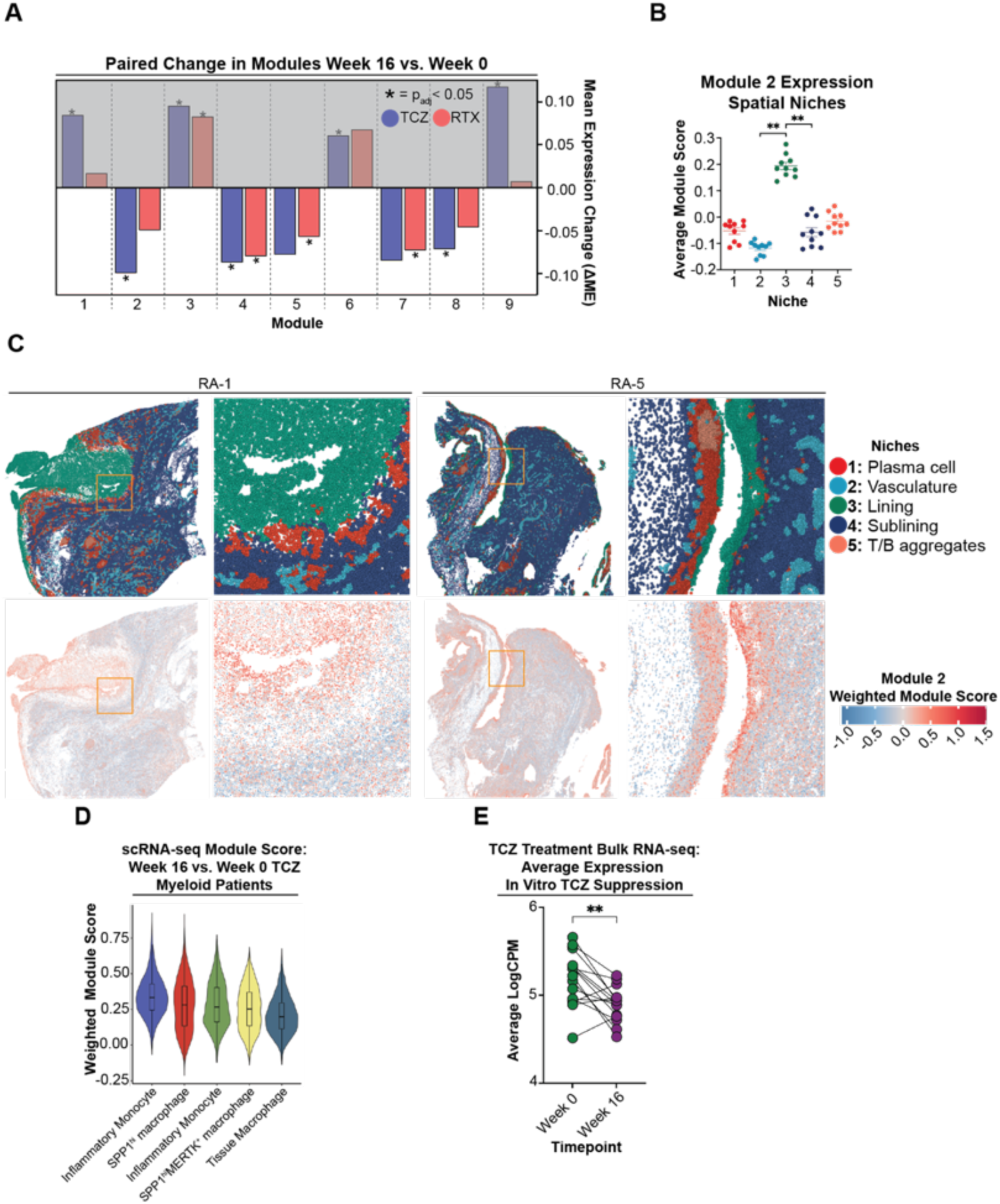
*SPP1*^hi^ macrophage targeting by IL-6 receptor inhibition in RA synovium. **A.** Paired analysis of WGCNA module eigengene changes (ΔME; W16 vs W0) in patients treated with tocilizumab or rituximab. Modules are separated into upregulated (top panel, shaded grey) and downregulated (bottom panel), with the latter highlighted as the drug-targeted modules of interest. **B**. Quantification of Module 2 enrichment score across spatial niches within individual patient tissue samples. Each point represents a niche score within a single sample; n = 10. Wilcoxon matched-pairs signed rank test: **Niche 3 vs. Niche 2 p = 0.002; **Niche 4 vs. Niche 3 p = 0.002. **C.** Spatial niche cluster plot (top panels) and spatial feature plot (bottom panels) showing the enrichment of Module 2 gene expression in a representative synovial tissue section. For spatial feature plot, Module scores were capped at the 99.5th percentile to improve contrast. **D.** Average expression score, calculated per cell type from synovial scRNA-seq data, for a gene module downregulated by tocilizumab in Myeloid-rich patients (derived from bulk RNA-seq). The plot displays the top 5 highest-scoring cell types: *IL1B*^+^*HBEGF*^+^ inflammatory monocytes/macrophages, *SPP1*^+^*CLEC5A*^+^ macrophages, *STAT1*^+^*CXCL10*^+^ inflammatory monocytes/macrophages, *MERTK*^+^*HBEGF*^+^ macrophages, and *MERTK*^+^*SELENOP*^+^ macrophages. **E.** Expression score of genes downregulated by tocilizumab in an in vitro *SPP1*^hi^ macrophage model, shown in patient synovial tissue pre-(Week 0) and post-(Week 16) tocilizumab treatment; n = 15; **Wilcoxon matched-pairs signed rank test p = 0.0067.

Comparing the bulk RNA-seq data from the Myeloid pathotype synovial samples before and after tocilizumab conditions to previously defined RA synovial cell type signatures, we found the top five cell types predicted to be downregulated by the treatment included two inflammatory monocytes states, two *SPP1*^hi^ macrophages subsets and a sublining tissue macrophage (**Figure 5D**). By contrast, the B cell depleting therapy rituximab was predicted to predominately downregulate two plasma cells, one B cell and two inflammatory monocyte subsets (**Supplementary Figure 5A**). These data further support prior analyses that the IL-6 receptor inhibitor tocilizumab effectively reduces myeloid populations in RA synovium (*35, 38*), while now demonstrating this includes *SPP1*^hi^ macrophages. Further, genes downregulated by tocilizumab in a bulk RNA-seq analysis of our in vitro generated *SPP1*^hi^ phenotype (macrophages co-cultured with synovial fibroblasts and TGF-β) showed a decrease in patient samples after treatment with tocilizumab (**Figure 5E, Supplementary Figure 5B**). These data suggest *SPP1*^hi^ macrophages may be a prominent target of the IL-6 receptor inhibitor tocilizumab in RA synovium.

## Discussion

Hyperplastic growth of synovial tissue and pannus formation are hallmarks of RA, yet the mechanisms responsible are poorly understood. Enabled by spatial transcriptomic profiling and reconstituted tissue remodeling assays, we have identified *SPP1*^hi^ macrophages and fibrin as key components in this process. Our findings support a model of de novo tissue generation on the fluid-interfacing surface using fibrin aggregates as a template. In the fibrin scaffolds, abundant *SPP1*^hi^ macrophages with fibrinolytic activity promote fibroblast proliferation and an activation state primed for nascent extracellular matrix production.

Prior findings of abundant *SPP1*^hi^ macrophages in RA synovium have been challenging to reconcile as these cells are established mediators of tissue fibrosis, yet RA is not a prototypical fibrotic disorder (*5, 7–11, 19*). Our transcriptome-wide comparisons substantiated that these cells resemble macrophages from fibrotic lung and liver disorders. Further, like their pro-fibrotic counterparts, synovial *SPP1*^hi^ macrophages anchor a niche with fibroblasts expressing a matrix remodeling program poised for collagen production. However, using spatial transcriptomics and birefringence histologic measures, we found these cells in lining niches devoid of the highly ordered collagen that marks fibrosis. We posit that this lack of overt fibrosis relates to the unique architecture of synovial lining, which is notably absent of an epithelial basement membrane (*41, 42*). Unlike the constrained, dense fibrotic aggregates that form within a rigid organ capsule, collagen produced near a compliant synovial fibroblast lining may drive tissue outgrowth. In this case, synovial *SPP1*^hi^ macrophages are more accurately termed “pro-generative” than pro-fibrotic. While synovial growth involving these macrophages may occur beneath the lining, additional observations suggest it also extends onto an exterior substrate.

Fibrin is an established pathologic feature in the inflamed RA joint, observed as dense, white, globular precipitates in the synovial fluid or at the tissue surface (**Figure 6A**) (*43–45*). Using three orthogonal and corroborative histochemical stains, we detected sizeable fibrin deposits adhered to the synovial lining, where fibrin and collagen fibers interdigitated at the interface. Based on this pattern and the established role of fibrin as a scaffold for collagen deposition in wound repair, we hypothesize fibrin serves as a substrate for pathologic growth of synovial tissue (**Figure 6B**). Accordingly, in an animal model of inflammatory arthritis, Sanchez-Pernaute and colleagues found that fibrin first accumulates in the synovial fluid and then adheres and disrupts the tissue lining. Synovial cells then migrate onto the fibrin and form a lining layer around the fibrin projection, thereby incorporating it as a nascent extension of the tissue (*46, 47*). In this model, fibroblasts were predicted as critical mediators (*46*), while our data now identify *SPP1*^hi^ macrophages as prominent components and define the fibroblast phenotype participating in this form of synovial expansion.

**Figure 6.**
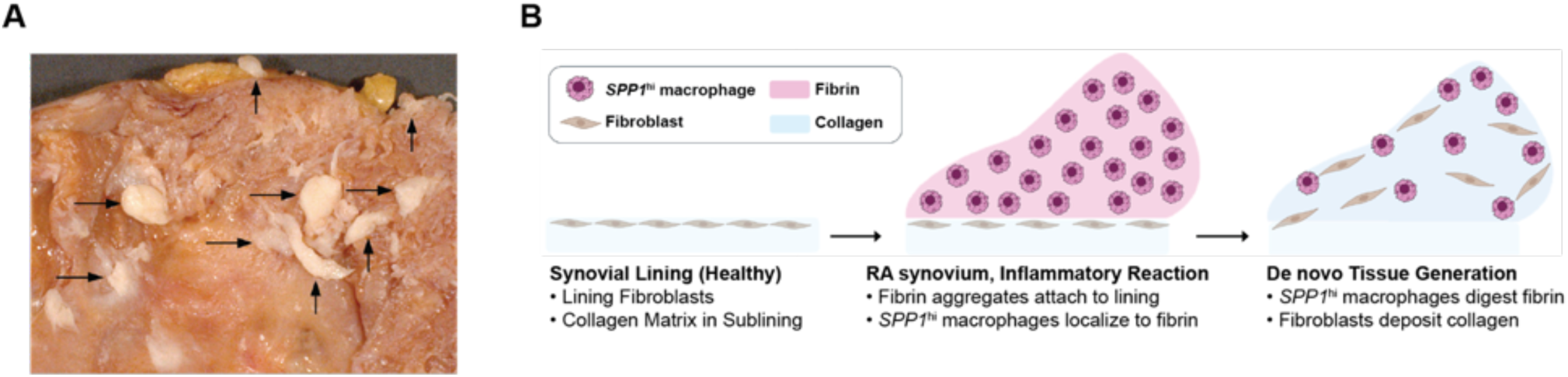
Model for de novo synovial tissue growth involving SPP1^hi^ macrophages and fibrin deposits. **A.** Gross image of the surface of the synovium from an RA patient showing the shaggy quality of synovial hyperplasia and numerous tan clots of fibrin adhering to the surface (arrows). **B.** Left: In healthy synovium, the synovial lining consists of lining fibroblasts that overlay the sublining collagen-based matrix (light blue) (white space represents synovial fluid and the joint cavity). Middle: In RA, fibrin exudates in the fluid cavity adhere to the lining surface and SPP1^hi^ macrophages localize to these fibrin matrices. Right: SPP1^hi^ macrophages digest the fibrin, where fibroblasts proliferate and produce matrix remodeling factors such as collagen.

By comparing the histochemical stains with spatial transcriptomics data, we detected a striking overlap of *SPP1*^hi^ macrophage aggregates and regions where fibrin dominated the extracellular matrix. Indeed, it was rare to observe these cells outside of fibrin scaffolds. *SPP1*^hi^ macrophages have also recently been found in synovial fibrin patches of newly onset juvenile idiopathic arthritis (*48*), collectively suggesting these cells are a principal feature of fibrin deposits in inflamed synovium. In a series of culture assays, we found that *SPP1*^hi^ macrophages efficiently degrade fibrin deposits and internalize fibrin fragments. As *SPP1*^hi^ macrophages have been found in tissues after acute injury (*49*), these cells likely function in a protective and restorative fashion during repair and homeostatic turnover to clear fibrin and other debris. However, in the RA joint, with unrelenting inflammation and excessive fibrin accumulation, we reason *SPP1*^hi^ macrophage activity is overwhelmed and persists en masse. As seen in spatial transcriptomics analysis, these macrophage co-localize with matrix remodeling primed fibroblasts which are now exposed to this substantial fibrin template.

The proposed function for fibrin as a template of pathologic synovial growth is distinct from its role in wound healing in key aspects. Most notably, in contrast to a wound, fibrin aggregates bound to the exterior of synovium provide a growth substrate where no tissue previously existed. Further, in a wound, the size and shape of the fibrin template is governed by both the boundary of the wound cavity and the new epithelial layer that grows over the top to resurface the wound (*50*). The joint cavity, in contrast, provides vast space for growth and an unrestrained structure, where nascent tissue morphology could instead be defined by the shape of the substrate. To this end, villous projections observed in RA synovium are morphologically consistent with fibrin exudates (**Figure 6A**) (*4*). Further, in one sample, we detected a particularly sizable fibrin attachment that extended outward more than 1 millimeter, roughly fifty-fold the thickness of a healthy lining (*41*). These features provide rationale for fibrin aggregates in the remarkably large and distinct patterns of synovial growth in RA.

Our findings inform a novel process relevant to RA treatment. Cohort biopsy studies have detailed most RA synovia contain *SPP1*^hi^ macrophages, with roughly a third dominated by these cells—deemed Myeloid-rich (*5, 19, 51*). Myeloid-rich tissues may represent a distinct RA endotype driven by divergent pathophysiologic mechanisms. Considering the overlap with *SPP1*^hi^ macrophages in our spatial data, we predict these samples also contain extensive fibrin deposits, and propose that fibrin is a key component to the pathologic response in Myeloid-rich tissues. Notably, patients with Myeloid-rich tissues have been found to respond particularly well to an IL-6 receptor inhibitor (*35, 52, 53*). We found in vivo targets of this response spatially localize to the synovial niche with *SPP1*^hi^ macrophages, proliferating fibroblasts and fibrin. Correspondingly, we and others have demonstrated IL-6 promotes *SPP1*^hi^ macrophage differentiation, along with MCSF and TGFβ (*8, 13, 49*). This suggests that *SPP1*^hi^ macrophages represent an unrecognized target of IL-6 pathway in RA, offering a candidate strategy for precision treatment in patients with Myeloid-rich tissues.

While our findings support a new model for RA-associated synovial tissue hyperplasia, they also highlight several limitations and avenues for future investigation. A key next step is to define the specific crosstalk factors mediating the interaction between *SPP1*^hi^ macrophages and fibroblasts, which could reveal novel therapeutic targets involved in pro-generative synovial remodeling. Furthermore, as our spatial analysis was performed on synovium from the knee, future studies should determine if this pathogenic niche is conserved across other joint types. Finally, the interplay between *SPP1*^hi^ macrophages and fibrin may represent a conserved pathogenic axis extending beyond RA. This may offer a new lens through which to view tissue remodeling in other inflammatory conditions such as lupus nephritis, where both components are implicated (*54, 55*), thereby broadening the potential impact of our findings.

## Acknowledgments

We thank HSS patients, orthopedic surgeons, rheumatologists, clinical research coordinators (particularly S. Cushing), the HSS Genomics core particularly Y. Chinenov, and D. Oliver for sequencing support. We thank W. Kang, N. Fan, and E. Rosiek at the MSK Molecular Cytology Core Facility (MCCF) for assistance with tissue sectioning, immunofluorescence staining, and imaging. The MSK MCCF is supported in part by the NIH/NCI Cancer Center Support Grant P30 CA008748. We are grateful to the NHGRI Impact of Genomic Variation on Function (IGFV) Consortium, which aims to understand how genome variation affects genome function and in turn phenotype. We acknowledge the R4RA Clinical Trial Network and C. Pitzalis for sequencing data generated in a synovial biopsy-driven trial. We acknowledge the the Accelerating Medicines Partnership Rheumatoid Arthritis and Systemic Lupus Erythematosus (AMP RA/SLE) Program for the large-scale synovial scRNA-seq dataset from patients with RA. AMP is a public-private partnership (AbbVie Inc., Arthritis Foundation, Bristol-Myers Squibb Company, Foundation for the National Institutes of Health, GlaxoSmithKline, Janssen Research and Development, LLC, Lupus Foundation of America, Lupus Research Alliance, Merck & Co., Inc., National Institute of Allergy and Infectious Diseases, National Institute of Arthritis and Musculoskeletal and Skin Diseases, Pfizer Inc., Rheumatology Research Foundation, Sanofi, and Takeda Pharmaceuticals International, Inc.) created to develop new ways of identifying and validating promising biological targets for diagnostics and drug development.

## Author contributions

Conceptualization: LTD, IM; Methodology: DO; Investigation: IM, RB, DR, EFD, HZ; Formal analysis: IM, HZ, AL; Data curation: HZ; Visualization: IM; Funding acquisition: LTD, KD, CSL, AR, TN; Project administration: MRF; Resources: DO, DR, EFD, SMG, MHS; Supervision: LTD, KD, CSL, AR, TN, FZ, KW; Writing – original draft: IM, LTD; Writing – review & editing: LTD, KD, CSL, AR, TN, MRF

## Data and materials availability

All data associated with this study are available or will be available in the paper, the supplementary materials, or through the Synapse data release. The Synapse release includes or will include raw and processed spatial transcriptomics data, raw and processed co-culture scRNA-seq data, raw bulk RNA-seq data from the polarization and co-culture studies, de-identified histology, birefringence, and immunofluorescence datasets, raw flow cytometry files with associated metadata and FlowJo workspaces, fibrinolysis assay source data, and sample-mapping and source-data inventory files. Raw co-culture scRNA-seq FASTQ files derived from human biospecimens are available through study-team-managed restricted access via the Synapse hub. Access requests may be submitted using the request form linked from the Synapse hub wiki, and approved users will be granted permission to download the files. RA synovium CITE-seq data used in this study from Zhang *et al.* (*5*) are publicly available on Synapse under the persistent identifier https://doi.org/10.7303/syn52297840.9. All code is available or will be available on Zenodo. All materials used or generated in this study are commercially available or will be supplied upon reasonable request.

## Methods

### 1. Experimental Design

The objective of this study was to investigate the cellular and molecular mechanisms of synovial tissue expansion in rheumatoid arthritis (RA). The study was designed as an observational and mechanistic analysis, combining high-plex spatial transcriptomics on human patient tissue with in-vitro functional assays using primary human cells.

RA synovial tissue for spatial transcriptomics analysis was obtained from 10 patients with seropositive RA (positive for RF and/or anti-CCP antibodies) who met ACR 1987 or ACR/EULAR 2010 diagnostic criteria. All patients were undergoing total knee arthroplasty (TKA) at Hospital for Special Surgery (approved by HSS IRB no. 2014-233). Enrollment complied with all relevant ethical regulations and informed consent was obtained from all participants. The patient cohort had an average age of 63.7 years (range: 46.6-76.5 years) and was 90% female. A diverse set of histological presentations were selected with the assistance of a clinical musculoskeletal pathologist (**Supplementary table 1B**).

Data inclusion and exclusion criteria were pre-established. For spatial transcriptomics analysis, cells were retained for analysis if they had at least 15 unique detected genes and 10 transcripts, and fell within 3 median absolute deviations (MADs) of the sample median for both gene and transcript counts. For scRNA-seq analysis, cells with more than 500 detected genes and less than 15% of transcript counts mapping to mitochondrial genes were included in downstream analysis.

For in-vitro functional assays, CD14+ monocytes were isolated from healthy donor leukopacks (New York Blood Center) and replicate cultures from these donors were subjected to different experimental conditions. The number of independent donors and experimental replicates (n) for each assay is specified in the corresponding figure legend.

The clinical musculoskeletal pathologists (D.R., E.F.D.) who performed the sample isolation and histological scoring were not blinded to the patients’ clinical information (including diagnosis, seropositivity, and other chart data), as this information was used to select a diverse set of histological presentations. Randomization was not applicable to the observational patient cohort analysis or in-vitro experiments, where paired donor cells were subjected to different treatment conditions.

### 2. Patient Sample Collection and Processing

#### a. RA patient synovial tissue samples selection

RA synovial tissue was obtained from a cohort of seropositive RA patients meeting ACR 1987 or ACR/EULAR 2010 diagnostic criteria undergoing total knee arthroplasty at Hospital for Special Surgery (approved by HSS IRB no. 2014-233). Enrollment complied with all relevant ethical regulations and informed consent was obtained from all participants.

All 10 patient tissues used in this study for spatial transcriptomics analysis were removed from patients with RA during a total knee arthroplasty (TKA) procedure performed at the Hospital for Special Surgery (IRB #2014-233). All patients selected met the RA diagnosis criteria set forth by the ACR in 1987 and/or the ACR+EULAR in 2010, and had a positive serum test for the presence of RF and/or anti-CCP antibodies. The average age of the patients was 63.7 years (range: 46.6-76.5 years); 90% were female. A diverse set of histological presentations were selected with the assistance of a clinical musculoskeletal pathologist (**Supplementary table 1B**).

#### b. Tissue sectioning

Formalin-fixed paraffin-embedded (FFPE) tissue blocks were sectioned to the required thickness using a microtome. Sections were then carefully mounted onto the appropriate slide and subsequently baked at 60°C to ensure proper adhesion, or dried for later use in spatial transcriptomics applications.

### 3. Xenium Spatial Transcriptomics Preparation

#### a. Xenium spatial transcriptomic analysis

Tissue sections mounted on the 10X Xenium slides were prepared according to the manufacturers protocol (10x Genomics Xenium In Situ Analysis User Guide, Document No. [Insert relevant document number or version]).

#### b. Custom 480-gene panel design

The panel includes all cluster-marker genes from the AMP RA/SLE Network’s 2023 study(*5*) for endothelial cells, myeloid cells, and stromal cells. We also included many genes that fell under the category of “general” already used by 10X Xenium in their human multi-tissue and cancer panel. Marker genes for SAMacs were also included(*8*), as well as genes known to be associated with fibrosis in previous studies(*56*).

### 4. Spatial Transcriptomics Data Analysis

#### a. Cell segmentation

Cells were segmented using Baysor (v0.7.0)(*20*), which applies a Bayesian mixture model to optimize cell boundaries by maximizing the joint likelihood of transcript positions and transcriptional composition. Negative control probes and transcripts with a Q-Score below 20 were excluded prior to segmentation. Baysor was run with the following key parameters: *min_molecules_per_cell_set* to median transcript count per cell based on the initial 10x nucleus expansion-based segmentation; *min_molecules_per_segment* set to half this median value; *n_clusters* set to 6; and *prior_segmentation_confidence* set to 0.3.

#### b. Xenium In Situ data preprocessing

Cells were filtered using both sample-specific and universal quality control thresholds. Specifically, cells retained for downstream analysis were required to have at least 15 unique detected genes and 10 transcripts, and to fall within 3 median absolute deviations (MADs) of the sample median for both gene and transcript counts. Gene counts for each cell were normalized to the total transcript count, multiplied by a scale factor of 10,000, and log-transformed after the addition of a pseudocount of 1.

#### c. Xenium In situ data integration

Following quality control, cells from individual samples were integrated using Seurat’s dictionary learning-based atomic sketch approach(*57*). For each tissue sample, 40,000 representative cells (“atoms”) were selected on the basis of leverage scores. Atoms were integrated with reciprocal PCA, with the number of principal components chosen at the elbow point and capturing 90% of total variance, and with *k.anchor* set to 10.

#### d. Xenium In Situ data clustering and annotation

Atoms were first annotated into six major lineages (fibroblast, myeloid, NK, T, B, and endothelial) using SingleR(*21*), with the AMP RA/SLE Network 2023 scRNA-seq dataset(*5*) as the reference, *de.method*=”wilcox”, and *de.n*=5. Subclustering within each lineage was per-formed using the Louvain algorithm across a range of resolutions(*58*), with the final resolution selected based on a manual inspection of marker genes. The subclusters were manually annotated based on the marker genes detected by Seurat’s *FindAllMarkers* with a Wilcoxon Rank Sum test. The full dataset was then projected into the atoms’ integrated space and annotated using the dictionary representation learned during integration, implemented by Seurat’s *ProjectIntegration* and *ProjectData* functions respectively(*57*).

#### e. Niche clustering analysis

Distinct spatial neighborhoods were determined based on cell type composition. Using the BuildNicheAssay function in Seurat, a “niche matrix” was constructed by calculating the count of each cell subtype of the 25 closest spatial neighbors of every cell. This niche matrix was then scaled and centered using the ScaleData function. K-means clustering was applied to the resulting niche matrix to partition all cells into k=5 distinct spatial niches.

#### f. Protein-Protein Interaction Network Analysis

To evaluate functional protein associations, the list of differentially expressed genes was submitted to the STRING database (version 12.0). The analysis was performed for Homo sapiens, and interactions were determined based on all active evidence channels. Interactions with at least a medium confidence score (≥ 0.4) were considered. The resulting network was assessed for overall interconnectedness, and the Protein-Protein Interaction (PPI) enrichment p-value was reported.

#### g. Gene set enrichment analysis (GSEA)

Gene Set Enrichment Analysis (GSEA) was performed using the desktop application. For the spatial analysis, a pre-ranked GSEA with phenotype permutation was used to determine if the 29 TCZ-responsive genes represented in the spatial transcriptomics panel were enriched in specific spatial niches. Separately, for the in vitro macrophage polarization experiment, a custom gene set of the 110 genes that were consistently downregulated in vivo by TCZ was tested to determine if it was also downregulated in the TCZ-treated macrophages.

#### h. Co-location quotients (CLQs) calculation

Pairwise CLQ values were computed to quantify spatial associations between cell types [6]. (*59*). For a given infiltrating cell type (B) and target cell type (A), the CLQ was defined as:

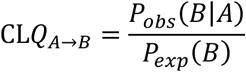

where the numerator represents the observed average proportion of an infiltrating cell type (B) in a *r* (50μm) radius neighborhood of the target cell type (A), and the denominator denotes the expected proportion of infiltrating cells given the overall sample composition. Empirical p-values were calculated by permuting cell positions under the assumption of spatial independence.

#### i. Niche differential gene expression (Niche-DE) analysis

Spatially modulated gene expression was assessed using Niche-DE, (*23*), which fits a negative binomial generalized linear model to test whether the expression of a gene in a given cell depends not only on its cell type identity but also on the density of neighboring cell types within its spatial niche. Genes were classified as “Niche-DE genes” if their expression in a specific cell type was significantly associated with the presence of other cell types, relative to the mean expression of that gene in the same cell type. The kernel bandwidth (*σ*), which defines the physical size of the local neighbourhood, was set to 175, 250, 300, corresponding to an average of 9, 20, and 30 spatial neighbors, respectively. To account for the skewed gene panel (larger number of representative genes for fibroblasts and myeloid cells) in our Xenium In Situ data, the γ parameter, a correction factor for cell type representation was reduced to 0.7 for fibroblasts and myeloid cells, while C, a threshold for the minimum number of cells required in a niche, was set to 150 and M, a filter for the minimum molecular count per gene, set to 50.

#### j. Image alignment (Xenium with histological sections)

To generate an outline of *SPP1*^hi^ macrophage-containing regions, the corresponding image (derived from Xenium-profiled tissues) was imported into GNU Image Manipulation Program (GIMP). *SPP1*^hi^ macrophage-containing regions were first isolated by using the “Select by Color” tool to remove background and *SPP1*^hi^ macrophage-devoid areas. Small internal voids within *SPP1*^hi^ macrophage regions were then removed using the “Remove Holes” tool, followed by the elimination of very small *SPP1*^hi^ macrophage-containing regions via the “Grow” and “Shrink” tools. A new transparent layer was subsequently added, and the refined selection was converted to a path. This path was then activated using the “Paths” tool, and stroke settings were configured to generate the outline. The original layers were then discarded, and the resulting outline was exported as a PNG image.

For spatial integration, this *SPP1*^hi^ macrophage outline was imported into Adobe Illustrator and precisely aligned with the image displaying both *SPP1*^hi^ macrophage-containing and-devoid regions. This composite image was then saved. The *SPP1*^hi^ macrophage layer within this composite was subsequently set to 50% transparency to facilitate its alignment with the H&E image generated from the corresponding Xenium-profiled tissue section. The H&E layer was then deleted.

### 5. Histology, Immunofluorescence, and Image Analysis

#### a. Analysis of histological images

Histological images were stained with Masson’s Trichrome, PTAH and H&E. Digital image analysis was performed on whole slide images of Masson’s Trichrome-stained tissue sections using QuPath (v0.6.0)(*60*). To quantify tissue composition, a supervised pixel classification workflow was developed to segment images into distinct tissue types based on color and texture.

On a single representative slide, a trained pathologist established the ground truth by drawing annotations around four distinct regions corresponding to two final classes:‘Fibrin+’ (representing pink-staining fibrin deposits) and’Fibrin-’ (representing blue-staining collagenous regions and areas of dense cellular infiltrate).

A Random Trees pixel classifier was trained on these annotated regions. The classifier was trained using a rich, multi-scale feature set designed to capture tissue characteristics at different magnifications. For each pixel, features included Red, Green, and Blue color intensity values, as well as image derivatives such as Gaussian blur, Hessian eigenvalues, and Structure Tensor eigenvalues calculated at three scales (sigmas of 0.5, 2.0, and 8.0).

The trained classifier was subsequently applied to the entire tissue area across all 10 slides in the cohort to generate a classification map for each sample. The total area (µm) for each class (’Fibrin+’ and’Fibrin-’) was then calculated from these maps. Classifier performance was validated by thorough visual inspection of the resulting color-coded classification maps to ensure accurate segmentation of the tissue components.

#### b. Quantitative analysis of overlap of birefringence with SPP1^hi^ macrophage containing regions

H&E stained sectioned that were previously labelled and processed for Xenium spatial transcriptomics analysis were imaged using circularly polarized light microscopy. Image channels were separated using QuPath software. The circular-polarized light (“blue”) channel was then processed in ImageJ (threshold: upper 15, lower 254) to generate a binary image. To enable background transparency, these binary images were subsequently imported into GNU Image Manipulation Program (GIMP), where an alpha channel was added and the white background removed. Birefringent regions were then pseudocolored for enhanced visualization. Finally, the processed images were aligned with their corresponding brightfield color images (captured during the same session) in Adobe Illustrator using standard alignment tools, leveraging their identical dimensions.

Spatial correlation between birefringence and specific cellular phenotypes was achieved through sequential image alignment in Adobe Illustrator. The previously generated birefringence region image, overlaid on its matched brightfield image (from the circular-polarized session), was precisely registered with the H&E image obtained from the Xenium profiled tissue section. This registration involved setting the brightfield image transparency to 50% and aligning it with the H&E image, followed by size adjustment and subsequent overlay onto the image representing *SPP1*^hi^ macrophage-containing regions. The intermediate brightfield layer was then removed. Prior to generating separate images, all layers were meticulously ensured to possess identical dimensions. To facilitate independent analysis while maintaining spatial registration, separate images were then generated for birefringence and *SPP1*^hi^ macrophage regions. These spatially registered images were then exported as PNGs on individual artboards to preserve identical dimensions and spatial coordinates. For quantitative colocalization, these images were imported into ImageJ. Binary masks were generated for both birefringence and *SPP1*^hi^ macrophage-containing regions (and *SPP1*^hi^ macrophage-devoid regions, using color thresholding). Finally, the “AND” function of the Image Calculator was applied to determine regions of overlap, and the number of colocalized pixels was quantified.

### 6. Cell Isolation and In Vitro Culture Experiments

#### a. In vitro macrophage polarization experiments

CD14+ monocytes were isolated from healthy donor leukopacks (New York Blood Center) using positive selection with anti-CD14 magnetic beads according to the manufacturer’s protocol (Miltenyi Biotec). Isolated cells were cryopreserved in liquid nitrogen in Cryostor (Biolife Solutions).

For experiments, cryopreserved CD14+ monocytes were thawed rapidly at 37°C and immediately transferred to pre-warmed RPMI 1640 (Corning) supplemented with 10% FBS (VWR, 76499-988), and 1% L-glutamine (Gibco, 25-030-081) (hereafter referred to as RPMI++). Following thawing, cells were washed twice with RPMI++. Cells were then seeded at a density of 1.0 x 10^6^ cells per mL, with 1 mL of cells per well, in 24-well tissue culture plates. Cells were either rested overnight in RPMI++ at 37°C in 5% CO2 or differentiated into macrophages by treatment with 50 ng/mL recombinant human M-CSF (Biolegend, 574804) for 24 hours.

After the initial resting or differentiation period, cells were treated with the following soluble factors at the indicated concentrations: M-CSF (50 ng/mL, Biolegend, 574804), IL-6 (20 ng/mL, Biolegend, 570804), TGF-β (20 ng/mL, Biolegend, 781804), TNF-α (10 ng/mL, Biolegend, 570102), and PGE2 (280 nM, Thermo, 233560100). Cells were incubated in media containing the respective soluble factors for the indicated time periods. Media was replenished every 48 hours by replacing half the volume with fresh RPMI++ containing 2X the initial concentration of soluble factors. Cells were harvested for downstream analysis or applications by gentle scraping of the well bottom and flushing of the media.

For the generation of macrophage conditioned media, macrophages were first washed three times with RPMI++. Cells were then counted and resuspended at a density of 2.0 x 10^6^ cells/mL. One mL of this cell suspension was added to each well of a 24-well tissue culture plate and cultured for 24 hours at 37°C in 5% CO2. Following the incubation, conditioned media was collected, centrifuged at 2,000 x g for 5 minutes to pellet cellular debris, and then filtered using a 0.22 μm syringe filter to ensure the complete removal of remaining cells and cellular components.

#### b. Co-culture with synovial fibroblasts

For co-culture experiments, human synovial fibroblasts (SFs) isolated from RA patients at passages 4-6 were seeded at a density of 7.5 x 10^4^ cells per well in the bottom chamber of 24-well plates containing 0.6 μm pore Transwell inserts (Corning). SFs were allowed to adhere overnight in RPMI++. The following day, the media in the bottom well was replaced with fresh RPMI++ containing the previously described soluble factors at their indicated concentrations. Where appropriate, tocilizumab (Selleck Chemicals, A2012) was added to the media at a concentration of 50 μg/mL. CD14+ monocytes, isolated and differentiated into macrophages by 24-hour M-CSF treatment as described in the “Cell Isolation and Culture” section, were then added to the upper chamber of the Transwell insert at a density of 6.0 x 10^5^ cells per well. The total volume in the upper chamber was 100 μL, and the bottom chamber contained 600 μL of media, as per manufacturer’s recommendations. Co-cultures were maintained for 72 hours at 37°C in 5% CO2. At the endpoint, macrophages in the upper chamber were harvested by gentle scraping with a P200 pipette tip followed by flushing. Synovial fibroblasts in the bottom chamber were harvested using TrypLE Express Enzyme (Gibco). For single-cell RNA sequencing (sc-RNAseq) experiments, harvested macrophages and fibroblasts were pooled at equal concentrations.

#### c. Fibroblast proliferation assay

Synovial fibroblasts were harvested and subsequently stained with 2.5 μM Cell Trace Violet (CTV) (Thermo Fisher Scientific, Catalog No. C34571) according to the manufacturer’s instructions. Stained fibroblasts were then seeded at a density of 2.0 x 10^4^ cells/mL, with 1 mL of cell suspension per well, into 24-well tissue culture plates containing αMEM++ (ThermoFisher, 41061029). Cells were allowed to adhere for 24 hours. Following the adhesion period, the αMEM++ was replaced with conditioned macrophage media, generated as described in the “In vitro *macrophage polarization experiments*” section. Where indicated, the conditioned media was supplemented with 3.6 μg/mL anti-Osteopontin (OPN) antibody (R&D Systems, Catalog No. MAB1433). Fibroblasts were incubated with the conditioned media for 72 hours. At the endpoint, fibroblasts were harvested using TrypLE Express Enzyme (Gibco, Thermo Fisher Scientific) and then stained for viability with eFluor780 Live/Dead Fixable Viability Dye (Thermo Fisher Scientific, Catalog No. 65-0865-14) according to the manufacturer’s protocol. Samples were then analyzed by flow cytometry. Flow cytometric data was analyzed using FlowJo software (BD Life Sciences).

### 7. Flow Cytometry

#### a. Viability Staining, and Surface Marker Staining

Macrophages were harvested from their respective in vitro culture or co-culture conditions and washed once with sterile phosphate-buffered saline (PBS). Cells were stained for 20 minutes on ice with either Live/Dead Fixable Aqua (Invitrogen, L34957) or eFluor 780 (Invitrogen, 65-0865-14) at a 1:1000 dilution in PBS. Following this, cells were washed once with FACS buffer (PBS supplemented with 5% fetal bovine serum and 2 mM EDTA). To minimize non-specific antibody binding, cells were incubated with a blocking solution composed of Fc Receptor Blocking Solution (Biolegend, 422302) at a 1:100 dilution and Monocyte Blocker (Biolegend, 426103) at a 1:20 dilution in FACS buffer for 30 minutes on ice. After the blocking step, cells were stained with a BV421-conjugated anti-CD14 antibody (Biolegend, 301830) at a 1:200 dilution in FACS buffer for 30 minutes on ice. Cells were then washed once with FACS buffer.

#### b. Fixation, Permeabilization, and Intracellular Staining

For intracellular staining, cells were fixed and permeabilized using a Cytofix/Cytoperm Fixation/Permeabilization Kit (BD, 554714) for 10 minutes as per the manufacturer’s protocol. Subsequently, cells were stained with a fluorescein (FITC)-conjugated anti-osteopontin (OPN) antibody (Bio-Techne, IC14331F) at a 1:100 dilution in permeabilization buffer for 30 minutes on ice. Following immunostaining, cells were washed twice with FACS buffer and resuspended in FACS buffer for data acquisition.

#### c. Data Acquisition and Analysis

Flow cytometry data was acquired on a BD FACSymphony A5 flow cytometer and analyzed using FlowJo software (BD). The gating strategy was as follows: Cells were first gated on forward scatter (FSC) and side scatter (SSC) to select the primary cell population, followed by sequential gating on FSC-A versus FSC-H and SSC-A versus SSC-H to exclude doublets. Live cells were identified by gating on the viability dye. The final macrophage population was identified as CD14-positive cells within the live, singlet gate. OPN expression was quantified as the median fluorescence intensity (MFI) of the FITC channel within this population.

### 8. RNA Sequencing and Bioinformatics

#### a. Bulk RNA-seq

Total RNA was extracted from tissue samples using the Qiagen RNEasy MicroKit, then reverse transcribed to cDNA. The cDNA libraries were prepared using the NEB Ultra II Directional RNA Library Prep kit, with the Poly A isolation module for mRNA enrichment. The libraries were sequenced on either an Illumina NovaSeq 6000 or Illumina NovaSeq X platform to generate approximately 40 million 150 bp paired-end reads per sample.

#### b. Weighted module generation

To refine the estimation of gene program activity, we implemented a weighted module score framework that incorporates the magnitude of differential expression. For each gene program (test set), control genes were sampled on the basis of average expression (stratified into 24 bins) to match the distribution of the test set. Weighted average expression values were then computed for both the test and control sets, with weights defined by the absolute log fold-change of each gene. The weighted module score was calculated as the difference between the weighted average expression of the test genes and that of the matched control set.

#### c. Similarity analysis (of scRNA-seq datasets)

A similarity matrix was constructed to quantitatively map and assess the relationship between myeloid cell subsets from the Fabre et al. (2023) dataset and the myeloid cell reference clusters from the Zhang et al. (2023) atlas. The Fabre et al. query cells, which contained pre-assigned cell type labels, were mapped onto the myeloid cell reference from the Zhang et al. study using the Symphony pipeline.

For cell label prediction, a k-nearest neighbors (k-NN) classifier was employed with a neighborhood size of k=5. This process leveraged the precomputed uniform manifold approximation and projection embeddings of the reference data to assign each Fabre et al. cell to its closest matching cluster.

A contingency table was then generated to enumerate the number of Fabre et al. cells from each subset that were assigned to each of the Zhang et al. myeloid reference clusters. To facilitate comparison and highlight similarities, the matrix was normalized both row-wise and column-wise using a Z-score transformation. The Zhang et al. cluster with the highest similarity score for each Fabre et al. subset (the column-wise maximum) was highlighted with an asterisk to denote the dominant mapping.

#### d. Correlation analysis

Statistical correlations were performed to assess the relationships between cell proportions and histological features. Specifically, we analyzed the correlation between the proportion of *SPP1*^hi^ macrophages and the percent area containing fibrin. We also correlated the proportion of total myeloid cells with the proportion of *SPP1*^hi^ macrophages within the myeloid population. This analysis was done using Prism software.

#### e. scRNA-seq analysis preprocessing

Cells with more than 500 detected genes and less than 15% of transcript counts mapping to mitochondrial genes were included in down-stream analysis. Gene counts were normalized using Seurat’s *SCTransform*(*61*), with mitochondrial mapping percentage regressed out. Doublets were identified using *DoubletFinder*(*62*), with *pK* optimized by the mean–variance normalized bimodality coefficient (BCmvn). Other parameters were set as follows: PCs = first 10 principal components, pN = 0.25, and nExp = the expected number of doublets (5% of total cells for 10,000 cells according to 10x) multiplied by the estimated heterotypic doublet rate.

#### f. scRNA-seq analysis annotation

Cells were first partitioned into macrophages and fibroblasts based on canonical marker expression. Each population was then annotated independently using Symphony (*63*), with reference mapping to the scRNA-seq atlas from (*5*), following the same procedures described therein.

### 9. In Vitro Functional Assays

#### a. In vitro clotting experiment

Macrophages were first polarized in vitro as described in the “In vitro macrophage polarization experiments” section. Following polarization, macrophages were harvested and resuspended at a concentration of 1.11×10^6^ cells/mL in pooled human plasma containing K2 EDTA (Innovative Research, IPLAWBK2E50ML).

A volume of 90 µL of this cell-plasma suspension was added to each well of a 96-well plate. Each experimental condition and donor was run in triplicate. To initiate clot formation, 10 µL of a 60 mM calcium chloride (CaCl2) solution was added to each well, resulting in a final CaCl2 concentration of 6 mM. The plates were then immediately transferred to a microplate reader pre-set to 37°C. The incubation was static, with no shaking. Clot formation was monitored over a 2-hour period by measuring turbidity at an absorbance of 450 nm every 5 minutes.

#### b. Fibrinolysis assay

Fibrin gels with a final fibrinogen concentration of 2.0 mg/mL were generated in a 24-well plate with a final volume of 250 µL per well. Two stock mixtures were prepared at half the final volume (125 µL). The fibrinogen mixture was prepared by combining fibrinogen (Sigma, 341576-100MG) with 10X PBS, 10X RPMI, and water to achieve final gel concentrations of 2.0 mg/mL for fibrinogen and 1X for both PBS and RPMI. The thrombin mixture was prepared by combining thrombin (Sigma, 605190-100U) with 10X PBS, 10X RPMI, and water to achieve a final gel concentration of 0.4 U/mL for thrombin and 1X for both PBS and RPMI. To form the gels, 125 µL of the thrombin mixture was gently pipetted into each well, followed by 125 µL of the fibrinogen mixture. The two solutions were mixed with a pipette tip to prevent bubble formation. The plates were then incubated at 37°C with 5% CO2 for 30 minutes to allow for polymerization.

Macrophages were cultured atop the newly formed fibrin gels under the same conditions described in the “In vitro *macrophage polarization experiments*” section. After a culture period of 6 days, the media and non-adherent cells were removed. The gels were then washed five times with sterile PBS. To assess the integrity of the fibrin gels, they were stained for 10 minutes by adding Coomassie Blue G-250 (Bio-Rad, 1610436). Following a second series of five washes with PBS, the gels were allowed to dry for approximately 15 minutes. The gels were then imaged using a Thermo Fisher Scientific iBright imaging system. During imaging, the plate was precisely positioned to ensure the well of interest was centered in the imaging field, avoiding any obstruction from the deep-well plate walls.

#### c. Fibrin phagocytosis assay

Macrophages were polarized in vitro for 72 hours with indicated soluble factors as described in the “*In vitro macrophage polarization experiments*” section. The cells were then harvested, counted, and resuspended at a concentration of 1.0×10^6^ cells/mL. A volume of 500 µL of this suspension was then added atop fluorescently labeled fibrin gels in a 48-well plate.

Fluorescent fibrin gels were manufactured in a 48-well plate as described previously, with the final gel volume modified to 150 µL per well. The total fibrinogen was comprised of unlabeled fibrinogen and fibrinogen-AF647 (Thermo Fisher Scientific, F35200), with 0.5% of the final concentration of fibrinogen substituted with the fluorescent conjugate. The cells were incubated atop the gels for 24 hours at 37°C with 5% CO2.

Following the incubation, cells were harvested by first incubating the gels with TrypLE Express (Corning, 1260401) for 30 minutes. The cells were then detached from the gel via repeated pipetting with PBS using a p1000 pipettor. The harvested cells were subsequently processed for viability staining, data acquisition, and analysis as previously described.

### 10. Weighted Gene Co-expression Network Analysis (WGCNA)

#### a. Data Input and Pre-processing

Gene expression data from the combined TCZ and RTX treatment datasets (“Merged” dataset), consisting of pre-normalized LogCPM values for protein-coding genes, were used as input. The ‘goodSamplesGenes’ function in the WGCNA R package (v1.73) (*64*) checked for and removed genes/samples with excessive missing values. We checked for sample outliers were identified via hierarchical clustering (‘hclust’, method=“average”), but none were removed.

#### b. Network Construction and Module Detection (Merged Dataset)

A co-expression network was constructed using the WGCNA R package. An unsigned network was built using ‘blockwiseModules’. A soft-thresholding power (β) of 8 was chosen using ‘pickSoftThreshold’ (R² > 0.85). The adjacency matrix was transformed into a Topological Overlap Matrix (TOM). Gene modules were identified using dynamic tree cutting (‘minModuleSize = 30’, ‘mergeCutHeight = 0.25’). Genes not assigned were placed in the grey module (Module 0). Module eigengenes (MEs), representing the first principal component of each module, were calculated (‘moduleEigengenes’). Gene module membership (kME) was calculated using ‘signedKMÈ.

#### c. Paired Analysis of Module Eigengene Changes (Fig 5A)

To assess treatment effects on Merged module activity, the change in module eigengenes (ΔME = ME_Week16 - ME_Week0) was calculated for each module within each patient having paired samples. The distribution of ΔME scores for each module was compared between the tocilizumab (TCZ) and rituximab (RTX) treatment groups using an unpaired t-test. P-values were adjusted across modules using the Benjamini-Hochberg method (FDR < 0.05).

#### d. Functional Annotation (ORA) of Modules

Functional enrichment analysis for Gene Ontology (GO) Biological Process terms, KEGG pathways, and Reactome pathways was performed on the gene lists comprising each Merged module (excluding grey). Over-representation analysis (ORA) was conducted using the‘clusterProfiler’ (v4.16.0) (*65*) and‘ReactomePÀ (v1.52.0) (*66*) R packages (‘enrichGÒ,’enrichKEGG‘,‘enrichPathway’ functions). Entrez IDs were used for enrichment, mapped from gene symbols using ‘org.Hs.eg.db’ (v3.21.0). The WGCNA gene universe (all genes passing initial filtering) was used as the background. P-values were adjusted using the Benjamini-Hochberg method (FDR < 0.05). Enrichment results were simplified using the ‘simplify’ function in ‘clusterProfiler’. Top significant terms were reported (see Supplementary Table 5A).

### 11. Scoring of In Vitro Gene Signatures on Bulk RNA-seq Data

A single gene signature representing the in vitro response to tocilizumab was defined based on differential expression analysis (limma) of bulk RNA-seq data from human CD14+ monocytes co-cultured. Specifically, top 100 downregulated genes (P < 0.05, unadjusted, sorted by logFC) in the “M_CSF+Fibs+TGFbeta+TCZ” condition compared to the “M_CSF+Fibs+TGFbeta” condition were selected.

The activity of this in vitro downregulation signature was quantified in the paired (Week 0 and Week 16) patient bulk RNA-seq samples from the TCZ treatment arm. For each patient sample, a score was calculated as the average logCPM of the genes constituting the signature. The change in signature score between Week 16 and Week 0 within patients was assessed for statistical significance using a Wilcoxon matched-pairs signed-rank test (Prism).

### 12. Statistical Analysis

Statistical methods for specific analyses are described in the corresponding figure legends and Materials and Methods sections. For in vitro experiments and paired tissue analyses, statistical significance was determined using two-sided paired t-tests or Wilcoxon matched-pairs signed-rank tests, as indicated in the figure legends. For unpaired comparisons, a one-sided Wilcoxon Rank Sum Test was used (Fig. 3C). Correlation analysis was performed using simple linear regression (Fig. 2F). Analyses were performed using Prism (GraphPad), R, or other software as specified. A p-value of less than 0.05 was considered statistically significant.

For spatial transcriptomics and scRNA-seq analyses, statistical methods included the Wilcoxon Rank Sum test for marker gene identification, permutation testing for Colocation Quotients (CLQs), and a negative binomial generalized linear model for Niche-DE analysis. For WGCNA differential module regulation was tested with an unpaired t-test. For high-plex genomic analyses (e.g., Niche-DE, WGCNA), p-values were adjusted for multiple comparisons using the Benjamini-Hochberg method, with a False Discovery Rate (FDR) < 0.05 considered significant.

## Supplementary Materials and Methods

Fig. S1. Spatial transcriptomics data processing, clustering, and niche analysis. Fig. S2. Histological validation of fibrin deposits and birefringence analysis. Fig. S3. Transcriptomic comparison to fibrosis and in vitro macrophage polarization. Fig. S4. Effect of fibrin on in vitro SPP1hi macrophage phenotype. Fig. S5. Transcriptomic analysis of macrophage:synovial fibroblast co-culture. Table S1. Xenium gene panel and patient histological metrics (related to Supp. Table 1A, 1B). Table S2. Spatial transcriptomics analysis data and gene lists (related to Supp. Table 1C-N). Table S3. Fibrin and birefringence quantification data (related to Supp. Table 2A, 2B). Table S4. In vitro polarization and scRNA-seq mapping data (related to Supp. Table 3A-C). Table S5. WGCNA module data (related to Supp. Table 5A).

---

**Supplementary Figure 1:**
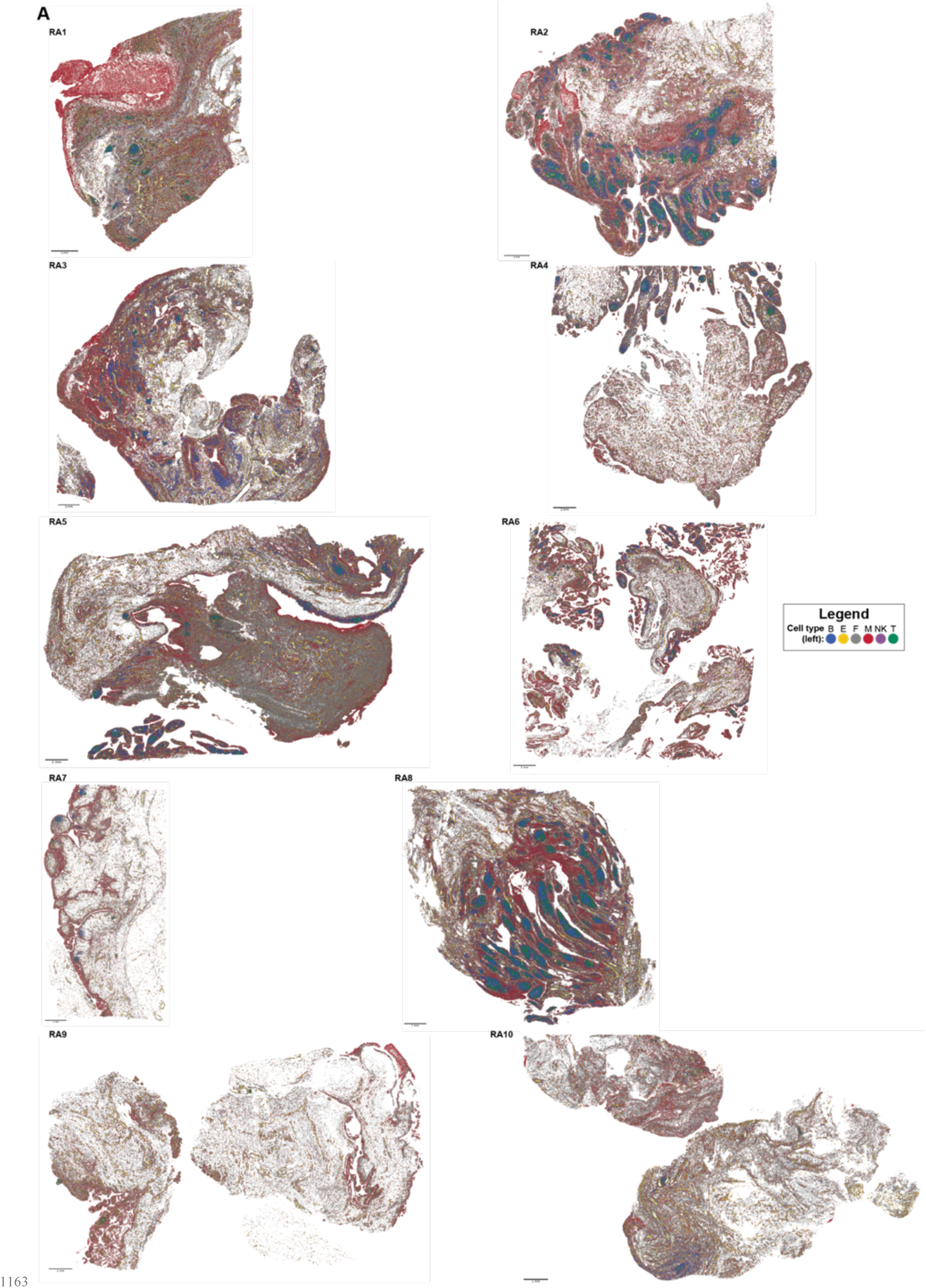
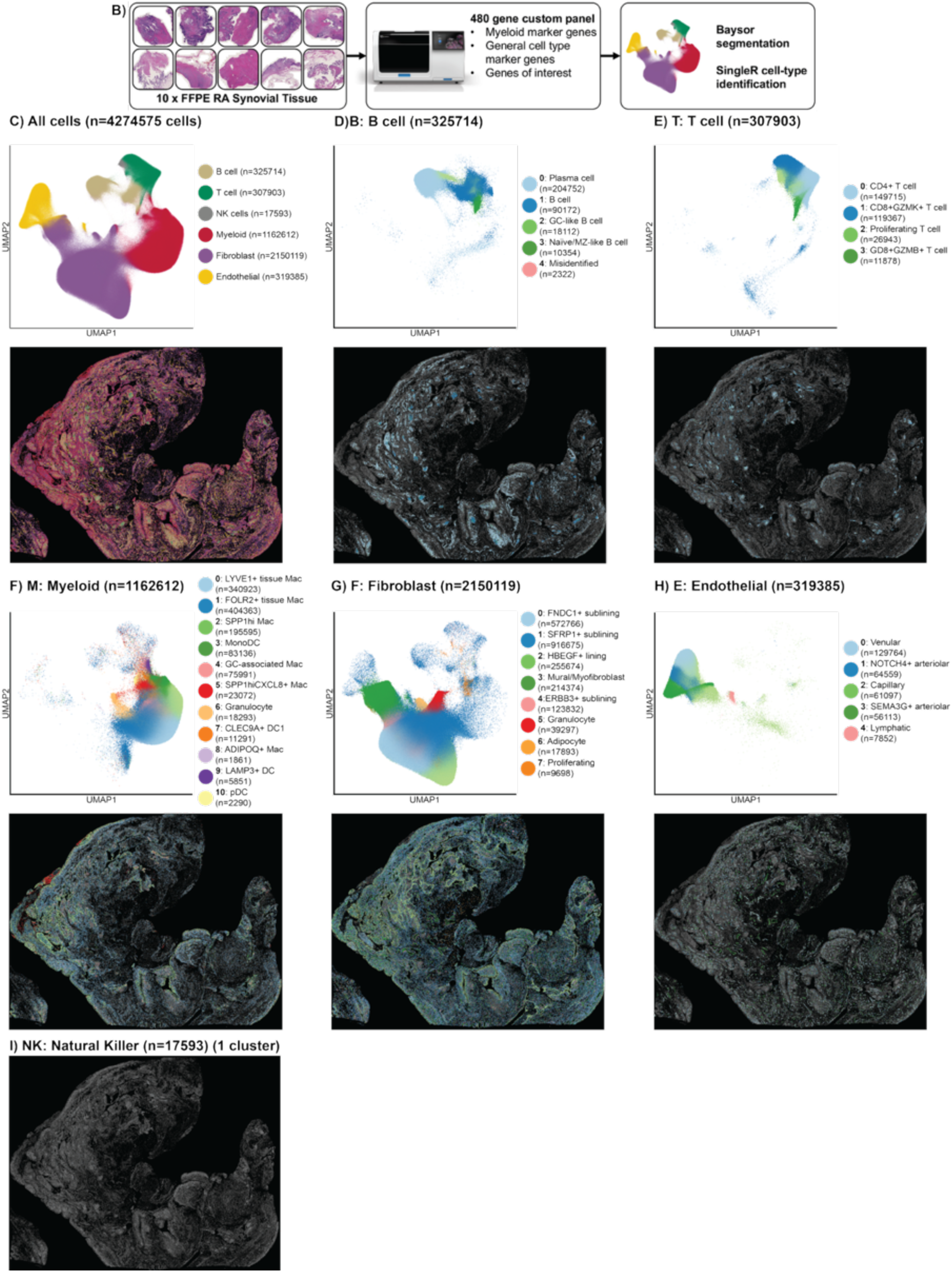
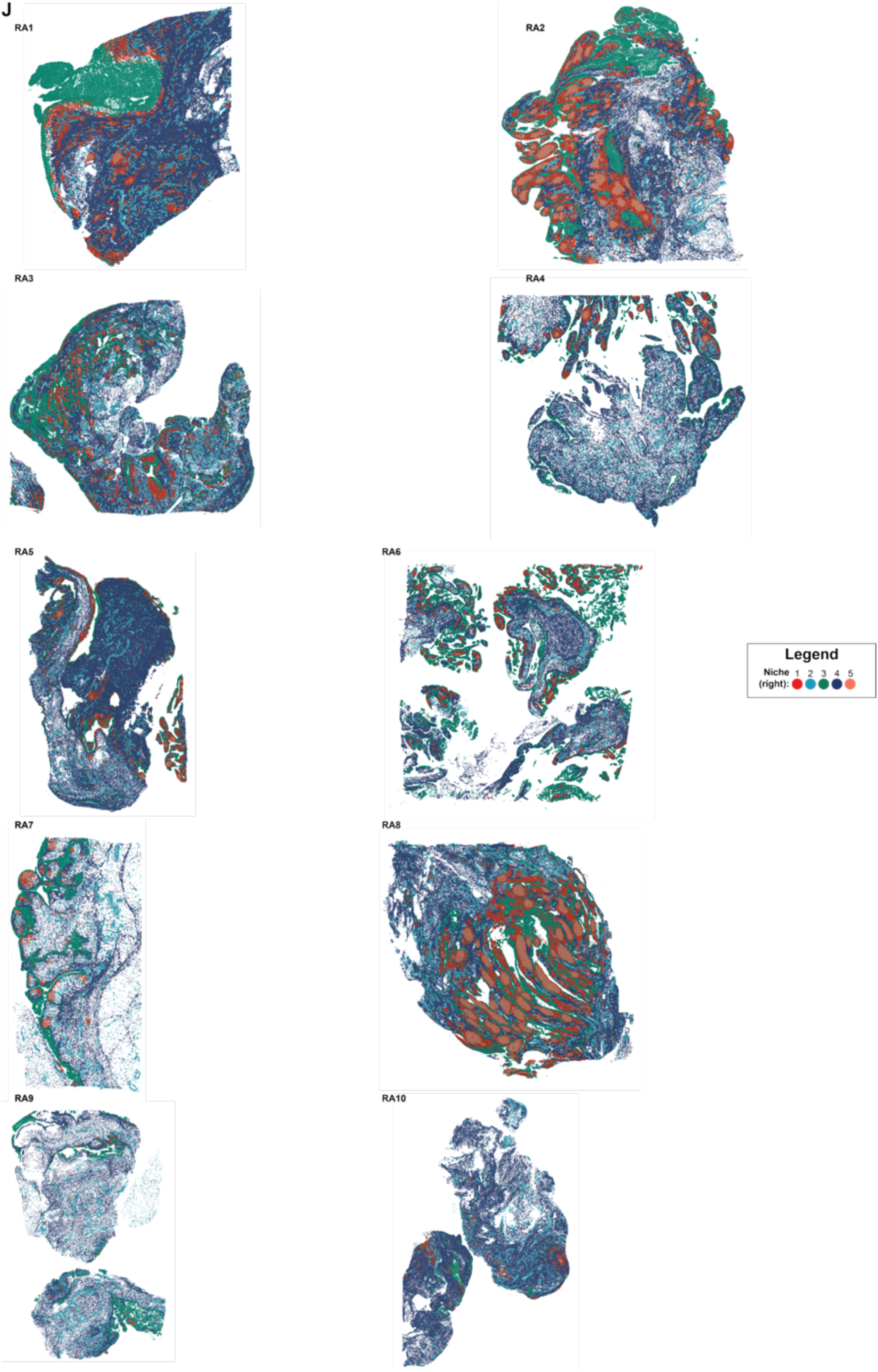
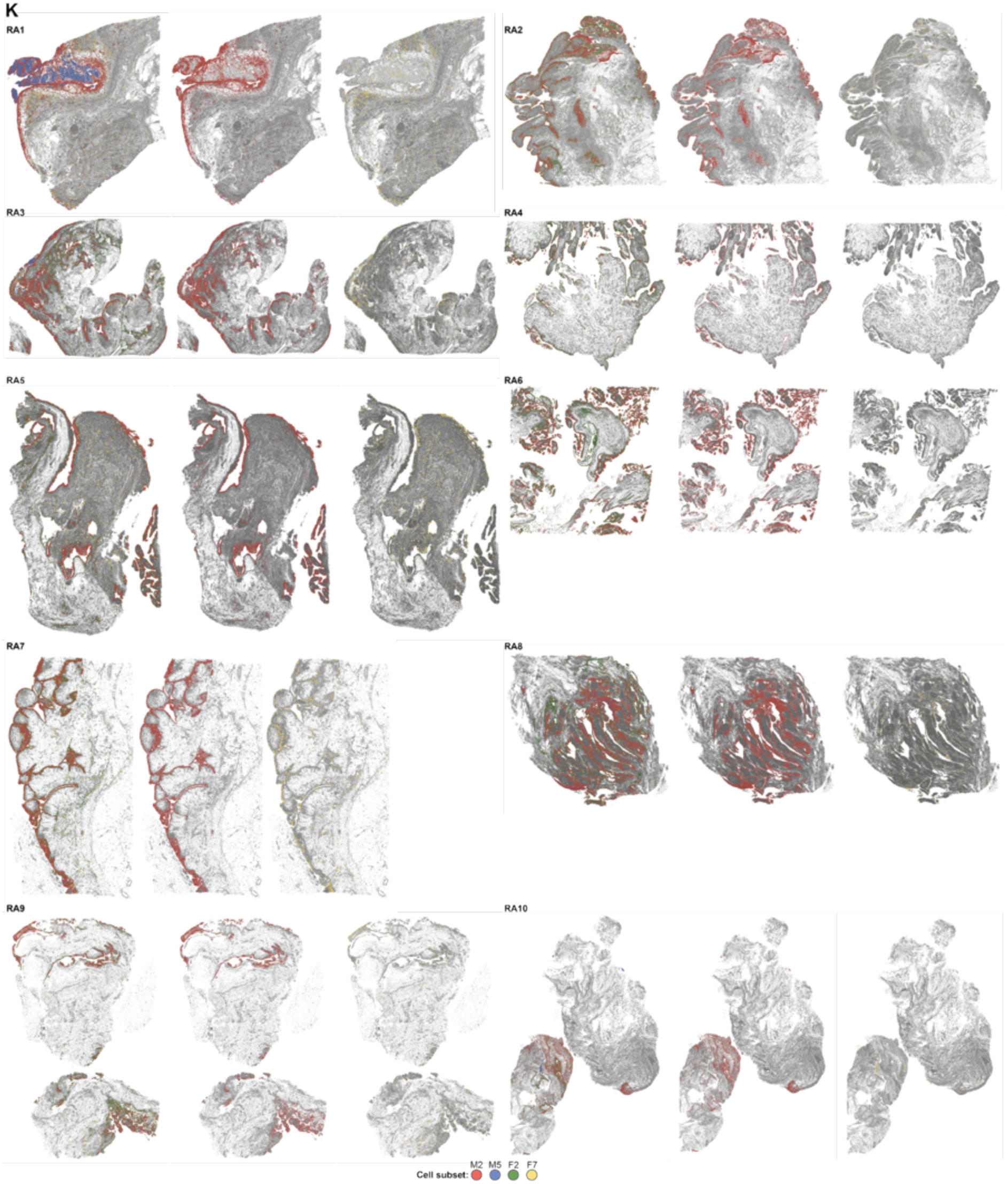
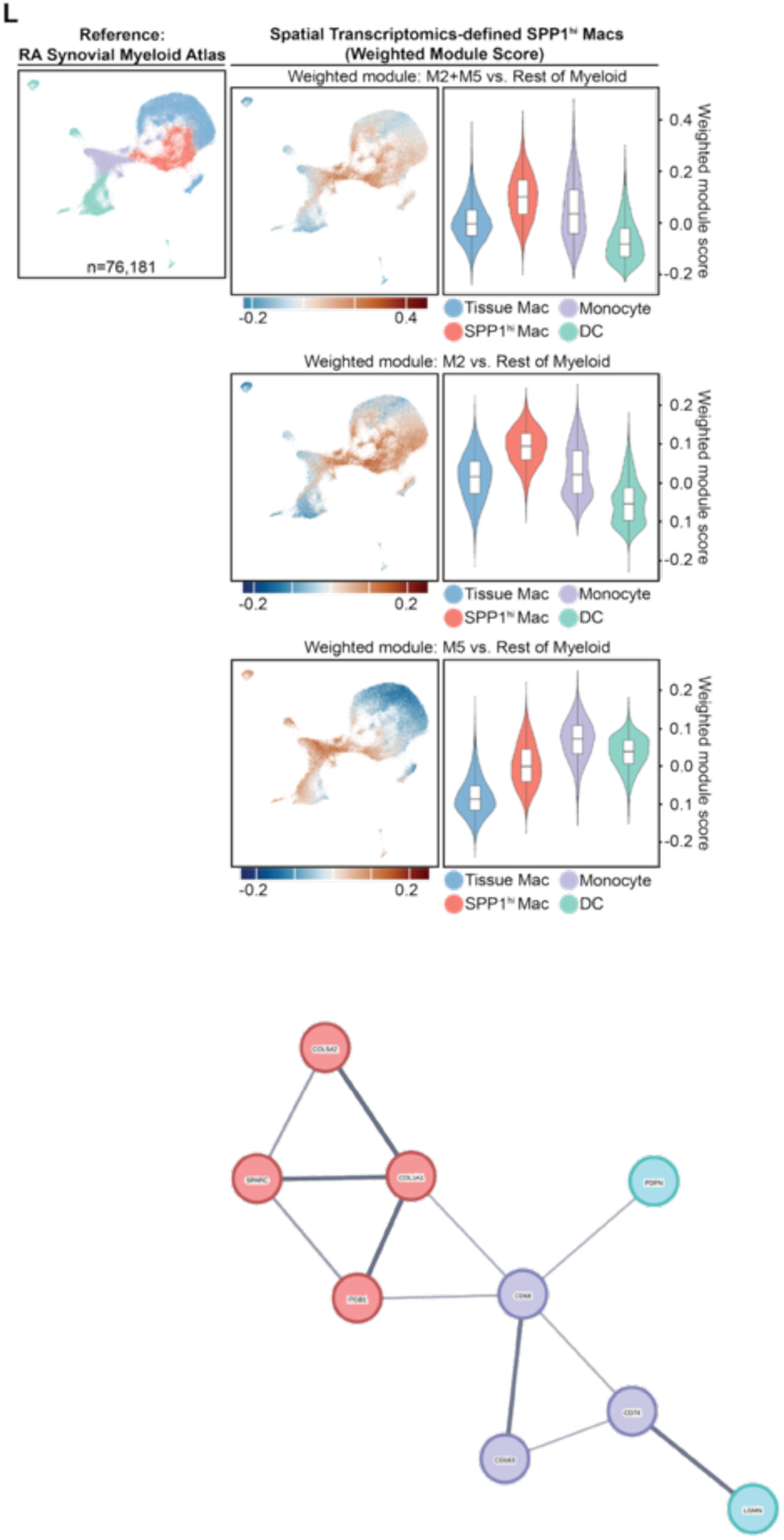
**A.**Representative images of each of the 10 RA patient synovial tissues profiled, with cells colored by broad cell lineage assignment. **B.** Diagram illustrating the workflow for Xenium spatial transcriptomics sample preparation, data acquisition, and analysis pipeline. **C-I.** Uniform Manifold Approximation and Projection (UMAP) embeddings and corresponding representative spatial images displaying cell subcluster assignments for all cells (**C**), B cells (**D**), T cells (**E**), myeloid cells (**F**), fibroblasts (**G**), endothelial cells (**H**), and NK cells (**I**); the NK cell UMAP is omitted as cells were maintained as a single cluster. **J.** Representative spatial transcriptomics images for each of the 10 tissues, with cells colored by niche assignment. **K.** Representative spatial images from each tissue highlighting the locations of *SPP1*^hi^ macrophages (M2, red), *SPP1*^hi^*CXCL8*^+^ macrophages (M5, blue), lining fibroblasts (F2, green), and proliferating fibroblasts (F7, yellow). **L.** UMAPs (left) and violin plots (right) displaying the expression score of weighted gene modules derived from spatial transcriptomics-defined *SPP1*^hi^ macrophage subsets (M2-specific, M5-specific, M2+M5 combined) calculated across cell types in an external scRNA-seq reference dataset.

**Supplementary Figure 2:**
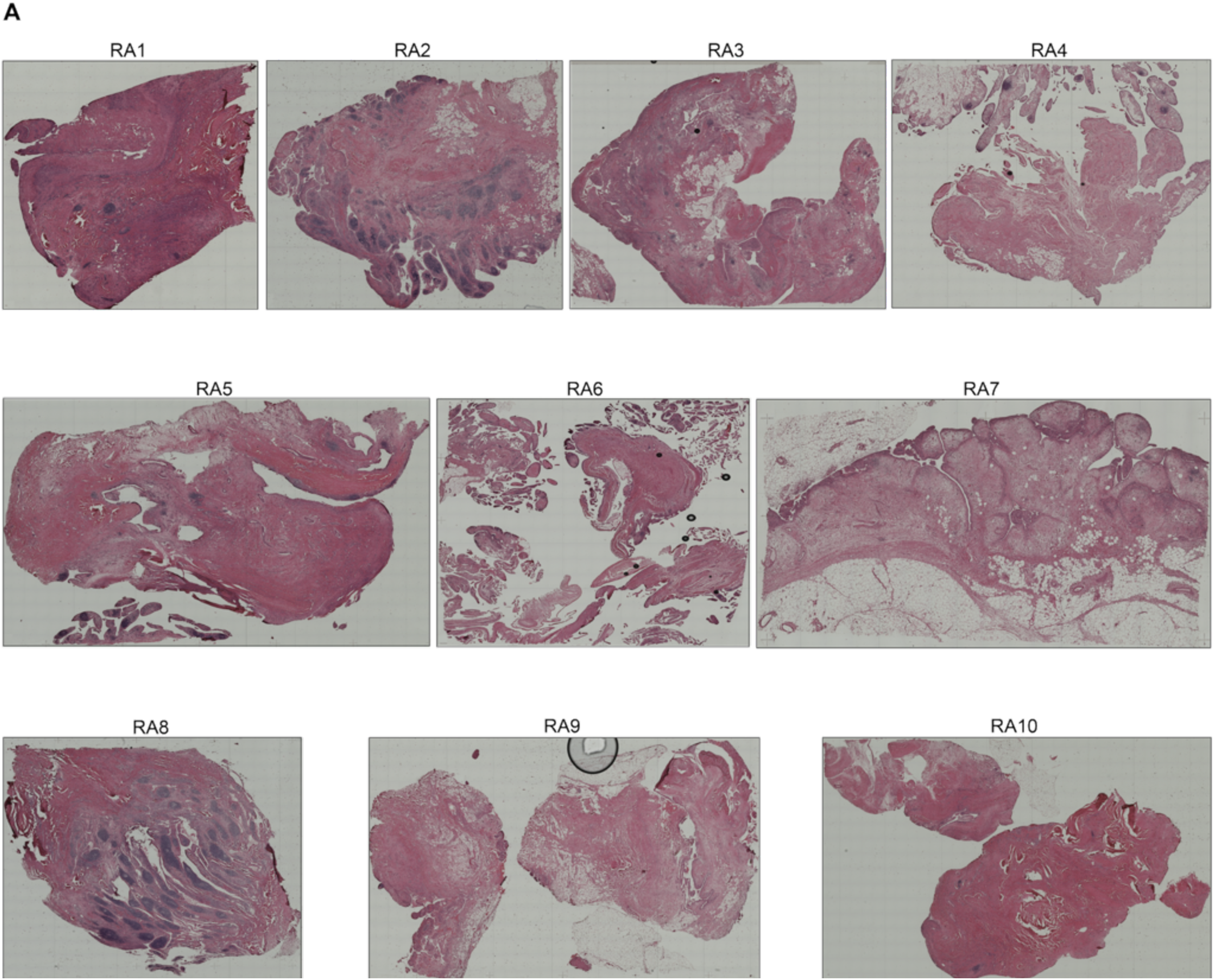
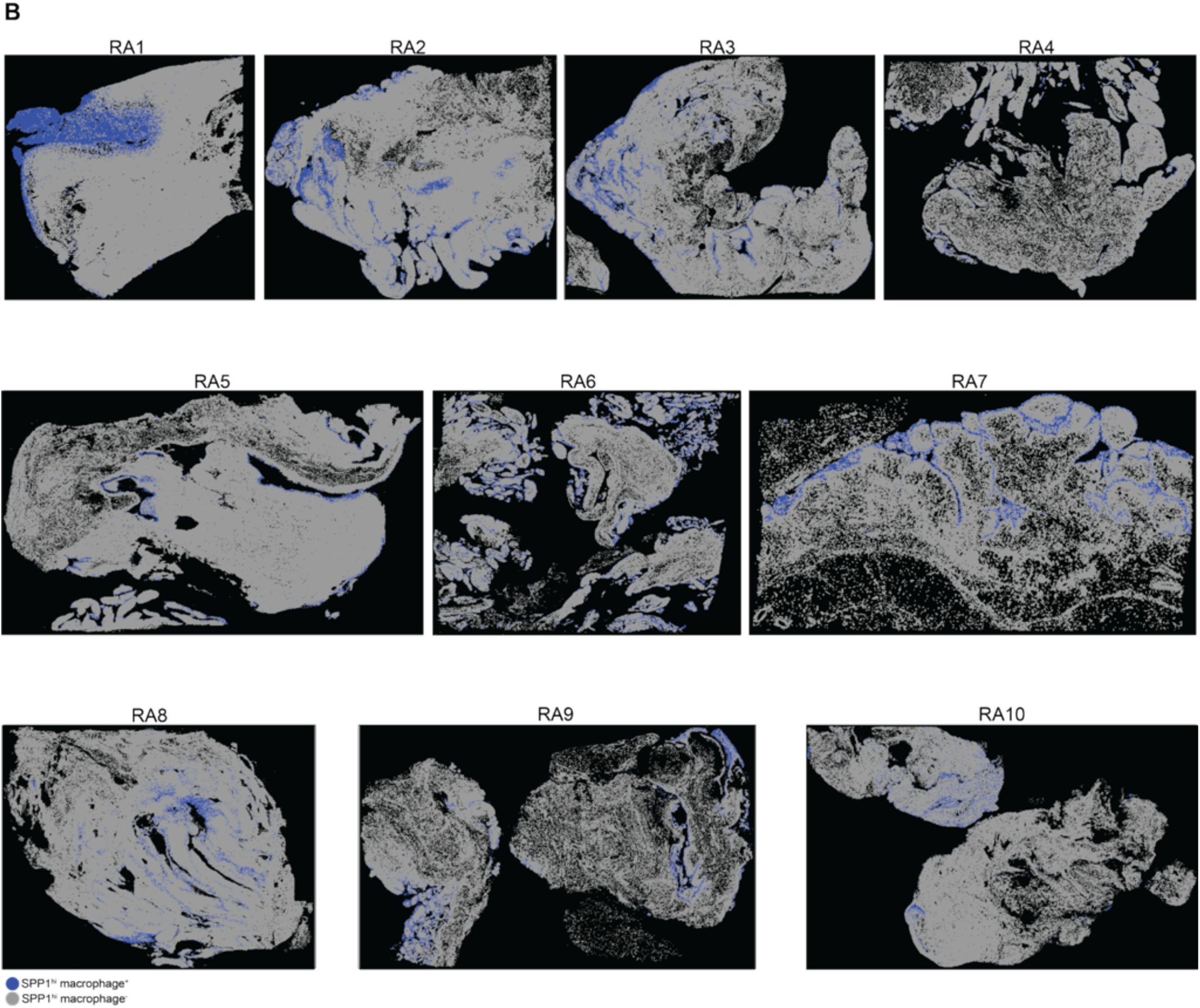
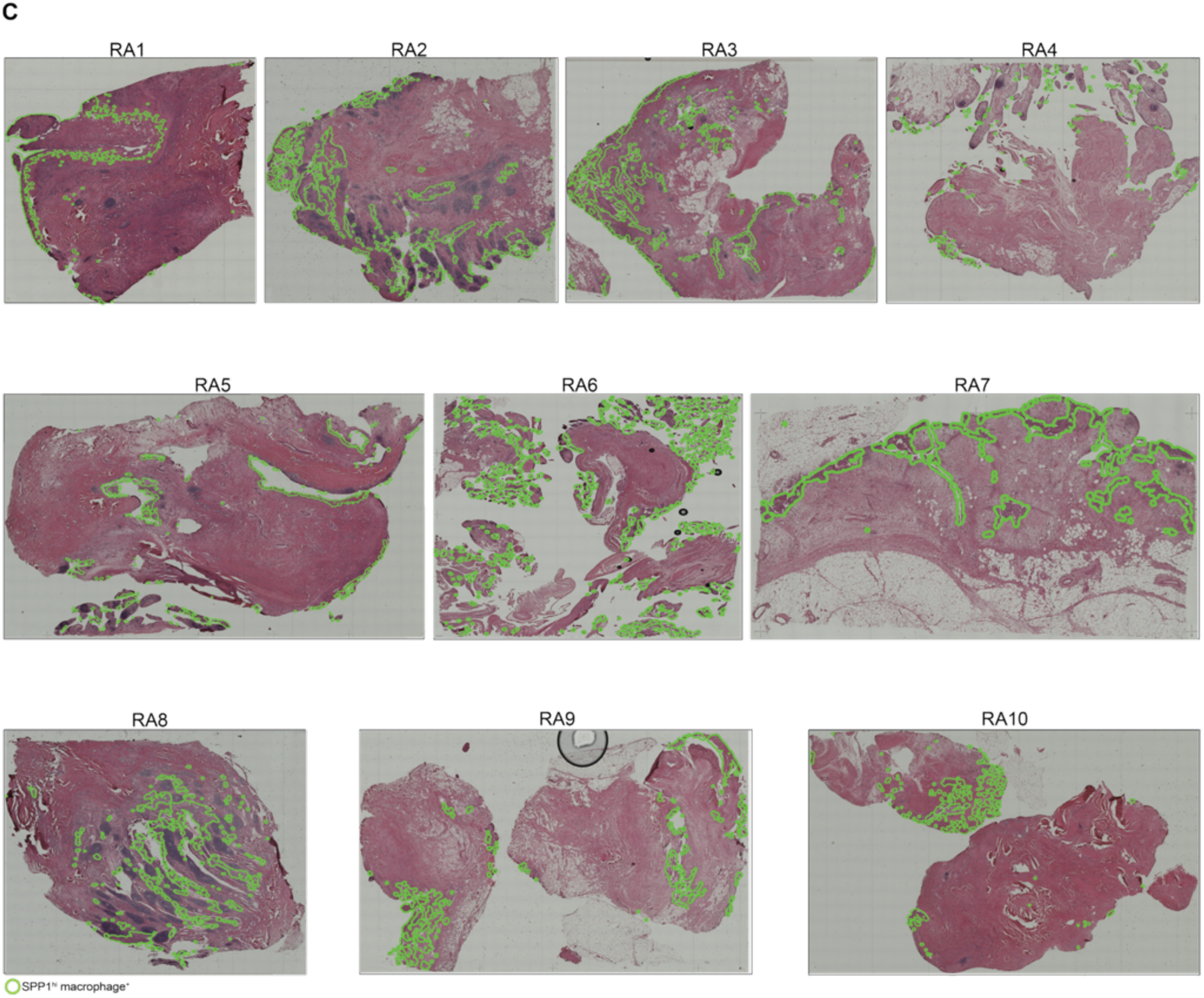
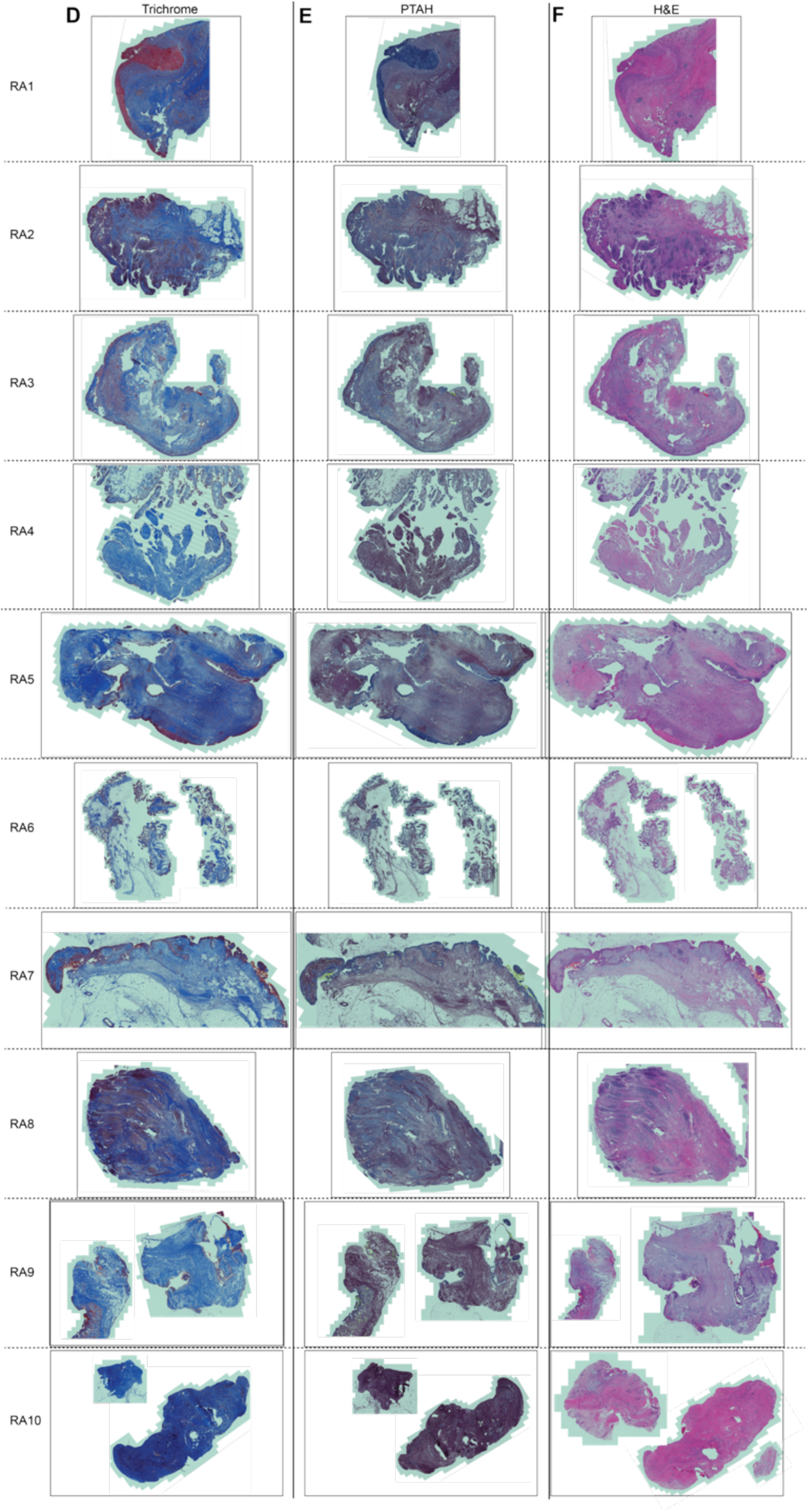
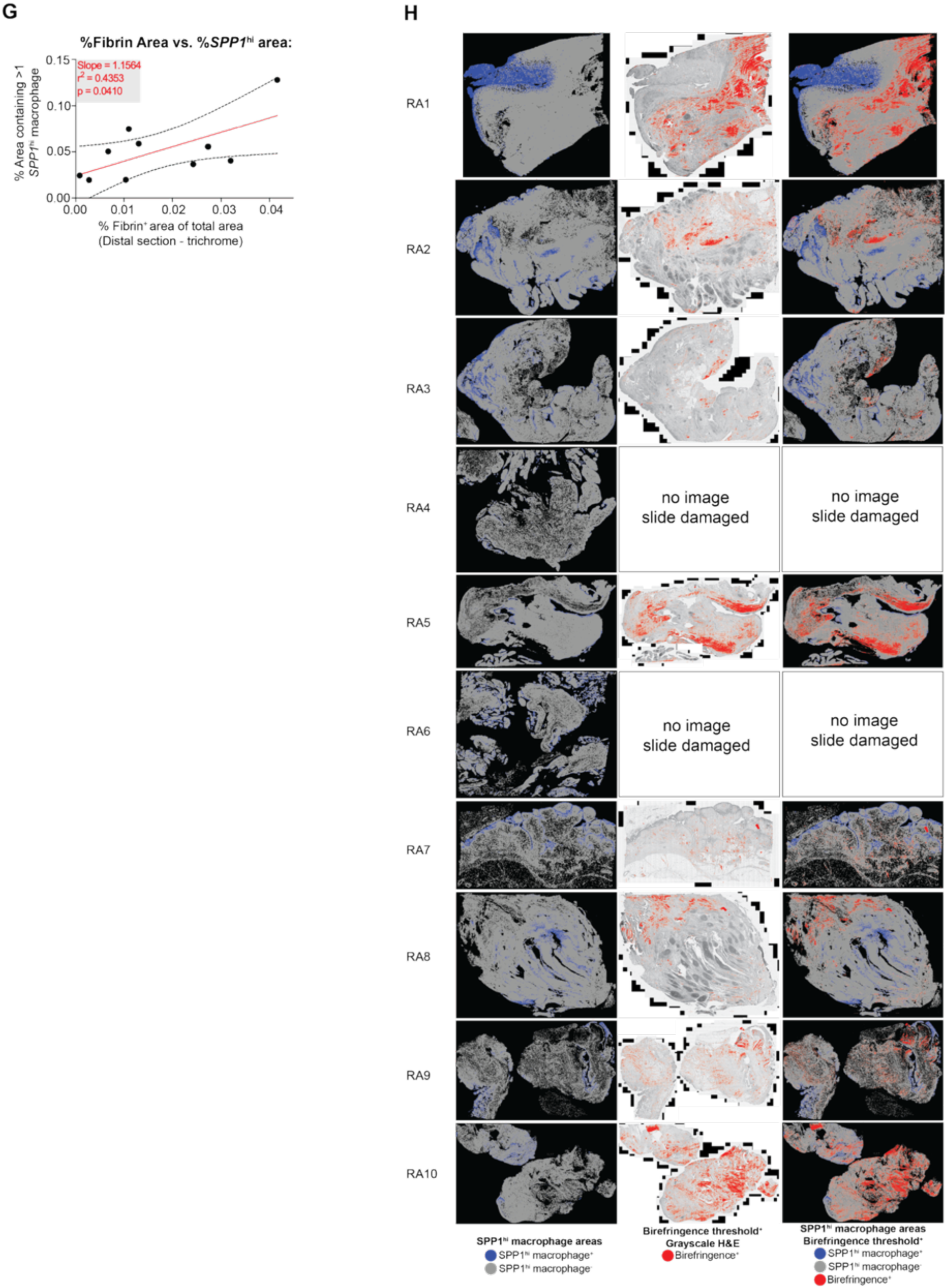
**A-C.** Visualizations of RA patient synovial tissue sections analyzed by spatial transcriptomics. (A) Hematoxylin and eosin (H&E) staining. **B.** Corresponding tissue regions computationally binarized based on the presence (blue) or absence (grey) of *SPP1*^hi^ macrophages. **C.** Outlines identifying *SPP1*^hi^ macrophage-containing regions (green) overlaid onto the H&E image from (**A**). **D-F.** Images of distal sections from the same tissue blocks stained with different histological dyes to visualize extracellular matrix components. (**D**) Masson’s Trichrome staining. (**E**) Phosphotungstic acid-hematoxylin (PTAH) staining. (**F**) H&E staining. **G.** Scatter plot correlating the percentage of tissue area identified as fibrin deposit (via object classification algorithm on Trichrome stain) with the total tissue area percentage occupied by *SPP1*^hi^ macrophages (M2+M5, determined by spatial transcriptomics). Includes linear regression fit and statistics. **H.** Additional representative related to the birefringence analysis using circularly polarized light microscopy, illustrating the low birefringence signals within and surrounding *SPP1*^hi^ macrophage-containing regions.

**Supplementary Figure 3:**
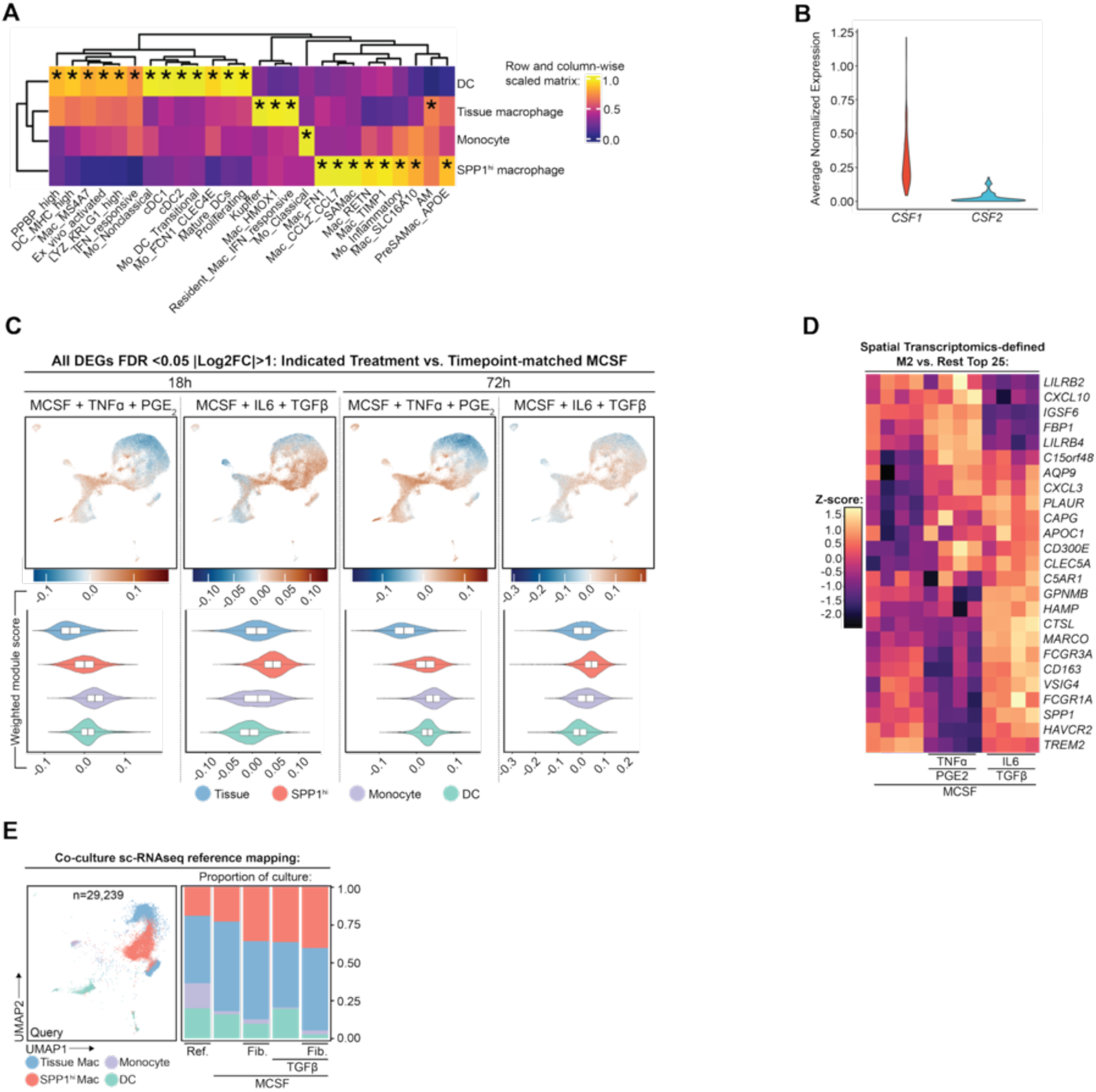
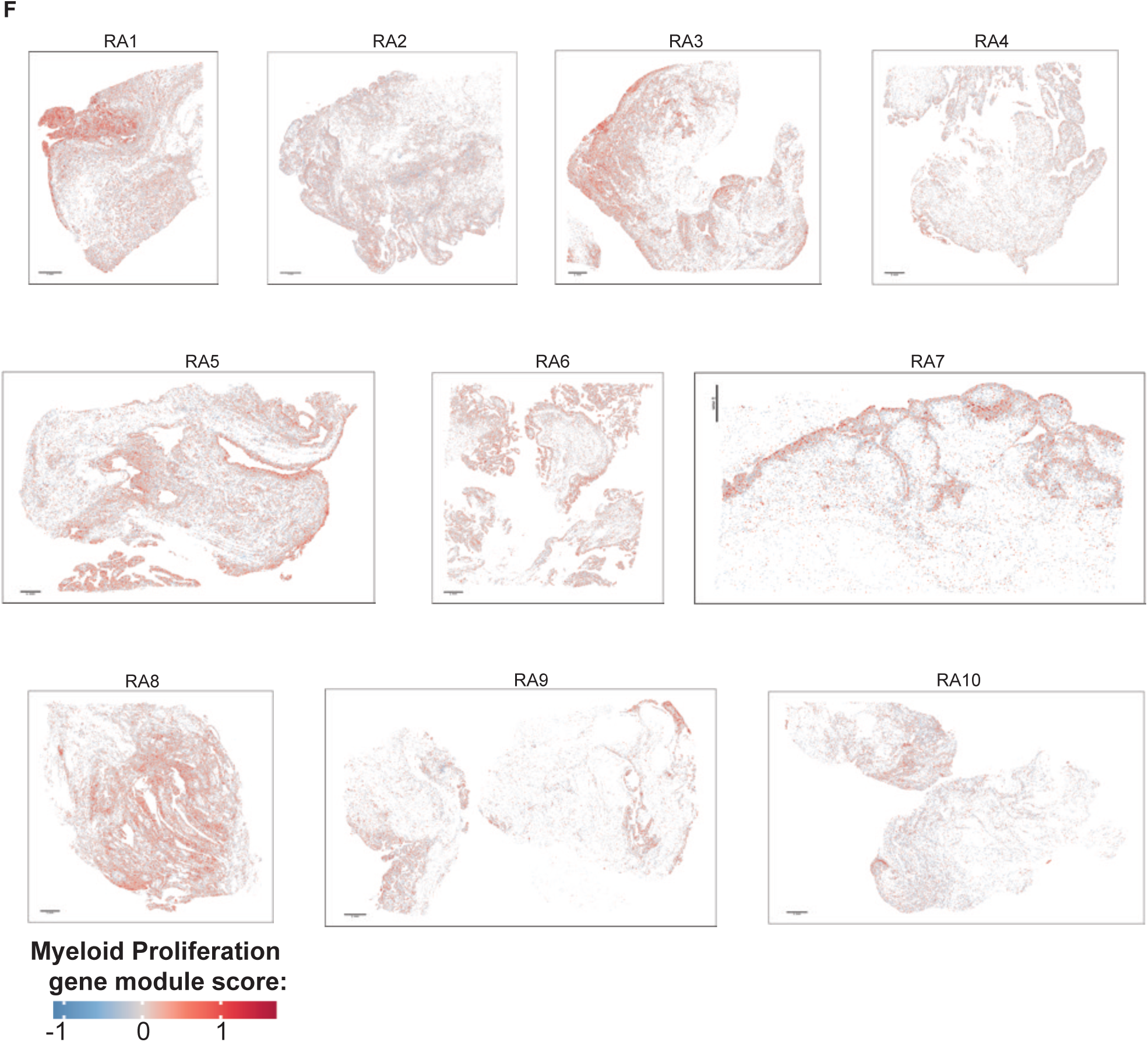
**A.**Similarity matrix comparing fibrotic lung/liver myeloid atlas to RA synovial myeloid atlas, showing highest similarity between lung/liver-derived fibrosis-associated, SPP1^hi^ SAMacs (fibrosis-associated macs) and RA synovial SPP1^hi^ macrophages, as well as similarity between expected cell subsets, such as classical monocytes and monocytes and various DC subsets. **B.** Violin plots showing higher expression of *CSF1* (M-CSF) in RA synovial tissue compared to *CSF2* (GM-CSF). **C.** Weighted module score corresponding to list of significant DEGs from bulk RNAseq data of CD14^+^ cells treated for 18h or 72h with M-CSF+TNF-ɑ+PGE2 (left side) or M-CSF+IL-6+TGF-β (right side) when compared to cells treated for 18h or 72h respectively with M-CSF alone; score of each cell in UMAP corresponds to legend under UMAP; violin plot shows weighted module score for each cell within indicated subset. **D.** Heatmap of gene expression in bulk RNAseq profiled CD14^+^ cells treated with the indicated soluble factors; gene list is top 25 DEGs based on Log2FC when spatial transcriptomics-defined SPP1^hi^ macrophages (M2) are compared with all other cells profiled by spatial transcriptomics; shows higher expression of M2 genes in cells treated with combination of M-CSF + IL-6 + TGF-β. **E.** sc-RNAseq data from macrophages (MACS sorted CD14^+^ cells treated with M-CSF for 24h) (co-)cultured as indicated for 72h; data is reference mapped back onto RA synovial myeloid atlas, with UMAP (left) and bar chart (right) showing highest proportion of SPP1^hi^ macrophages from culture output when cells cultured with M-CSF + TGF-β in presence of synovial fibroblasts. **F.** Spatial transcriptomics images depicting spatial distribution of expression of proliferation gene module in myeloid cells; this program overlaps with fibrin deposits and SPP1^hi^ macrophage containing-regions – this can be observed by cross-referencing these images with those in other figures.

**Supplementary Figure 4:**
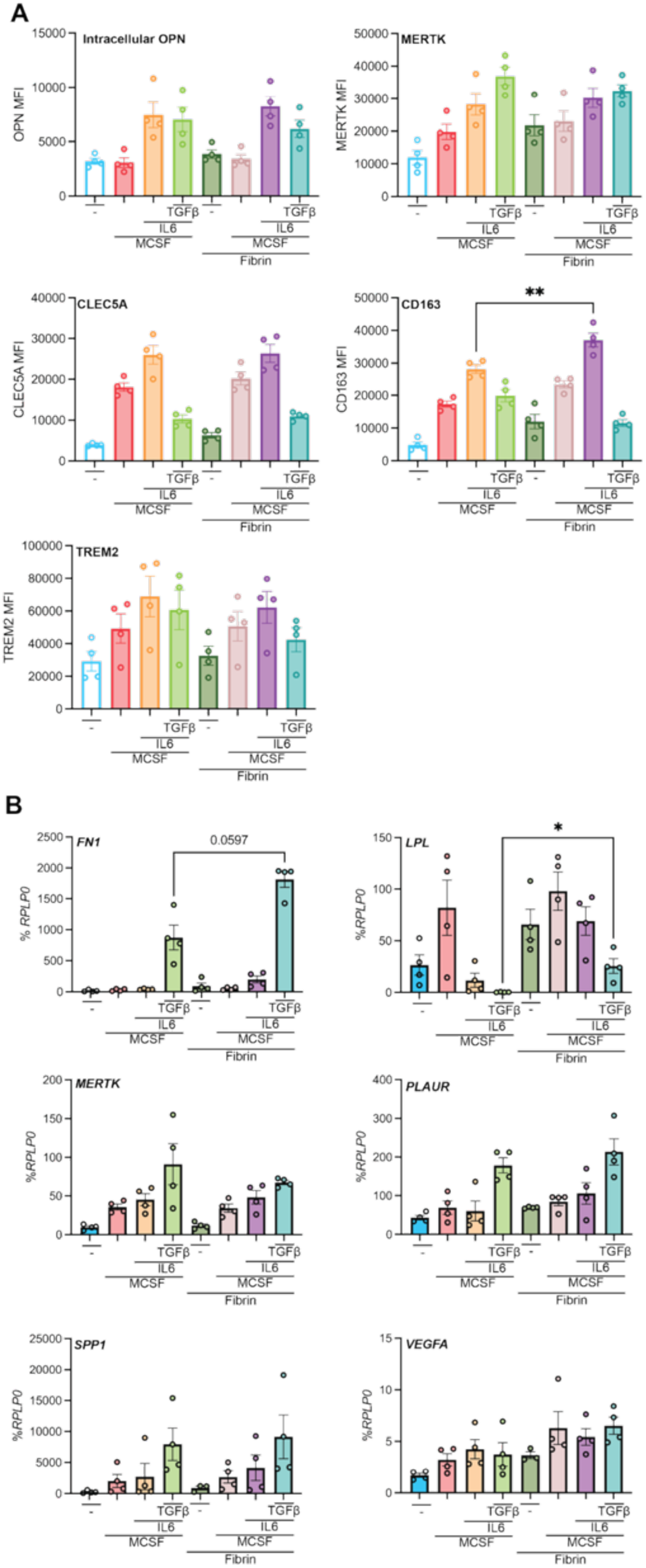
**A-B.**Flow cytometry (**A**) and RT-qPCR (**B**) of CD14^+^ cells cultured with indicated combinations of soluble factors with or without fibrin gel.

**Supplementary Figure 5:**
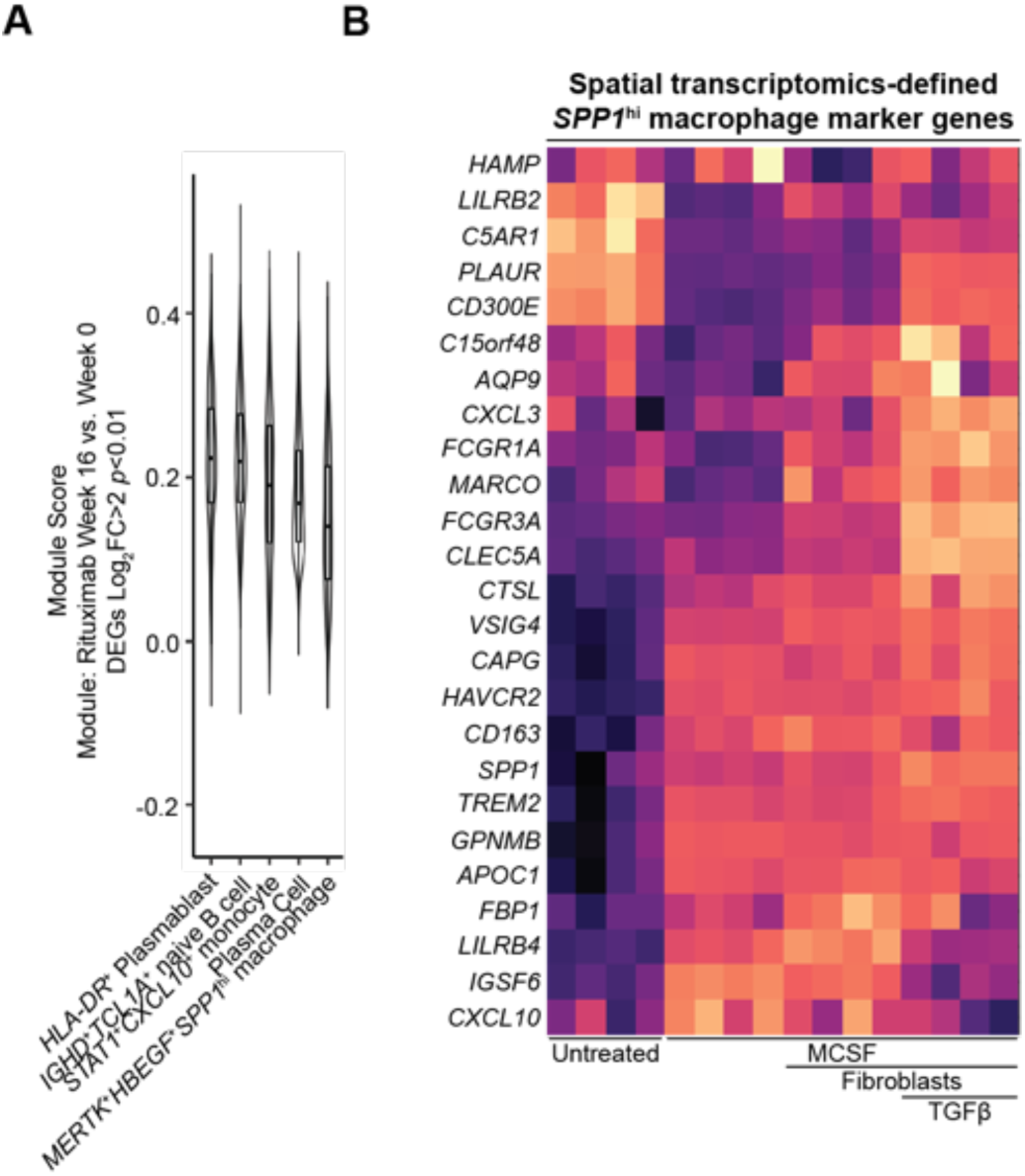
**A.**Violin plot showing weighted module score of scRNA-seq defined RA synovial cells. Module is based on bulk RNA-seq of patients treated with rituximab for 16 week, week 16 vs 0 DEGs with Log2FC > 2, *p* < 0.05. **B.** Heatmap of bulk RNA-seq data from macrophage:synovial fibroblast co-culture showing expression of top 25 spatial-transcriptomics defined *SPP1*^hi^ macrophage marker genes; the highest expression is observed when macrophages are co-cultured with synovial fibroblasts in the presence of TGF-β.

